# Uncovering Non-Monotonic Antagonistic and Synergistic Combinations (UNMASC), a robust method with applications to T cell differentiation *in vitro*

**DOI:** 10.1101/2025.09.07.674767

**Authors:** Geneviève Bistodeau-Gagnon, Yale S. Michaels, Morgan Craig

## Abstract

Induced pluripotent stem cells (iPSCs) have been proposed as an alternative T cell source for CAR-T cell therapies, as they can be differentiated and matured into T cells *in vitro* using cytokines. These assays benefit from computational and mathematical models to design appropriate experimental protocols. However, models are limited by typical monotonic dose-responses, preventing them from being used for cytokine effects that are largely non-monotonic. To address this shortcoming, we developed Uncovering Non-Monotonic Antagonistic and Synergistic Combinations (UNMASC), a novel mathematical model describing non-monotonic dose-response surfaces of cytokine interactions that distinguishes synergy of efficacy and potency. We showed that our approach successfully recapitulates non-monotonic observed dose-response surfaces characterizing *in vitro* T cell progenitor differentiation. Our results highlighted cytokine combinations with antagonistic effects alone but synergistic in combination, particularly IL3 and IL7. Together, UNMASC accelerates the efficient cell generation assays and extends drug interaction surfaces to a broader range of dose-responses.

## INTRODUCTION

Chimeric antigen receptor (CAR)-T cell therapy has shown great promise in treating blood malignancies^1-5^. Unfortunately, its use is limited by the need to harvest a patient’s T cells, which are largely suppressed due to cytotoxic chemotherapies^6-8^, and are cumbersome to engineer on a one-off autologous basis. Induced pluripotent stem cells (iPSCs) have thus been proposed as an alternative T cell source^6,9,10^. To generate T cells, iPSCs must first be differentiated into hematopoietic stem and progenitor cells (HSPCs) before T cell progenitor differentiation and maturation^6,9^. However, stimulation regimes yielding the highest number of mature T cells are difficult to establish due to the complex dose-responses cytokines often exhibit, many of which are non-monotonic.

That various cytokines, including those involved in T cell progenitor differentiation and maturation^6^, exhibit biphasic dose-response relationships leads to a poor understanding of their mechanisms of action. This limits their broad use as immunotherapies^11^. Further, drug-induced dose-response relationships are often described using saturating (i.e., “S”-shaped) models. However, similar non-monotonic dose-responses have been found in apoptotic^12^ and anti-angiogenic cancer drugs^13^, among others. Numerous biological processes, including endocrine and cytokine signals, also exhibit non-monotonic doses-responses. These include lymphocyte activation and proliferation which have both been shown to display stimulatory effects at low doses and inhibitory effects at high doses of cytokines and drugs^14^. Similar bell-shaped curves also describe dose-responses characteristic of interleukin (IL), including IL-1^15-17^, IL-6^17^, IL-11^18^, tumour necrosis factor-alpha (TNFa)^15,17,19^, and interferon-gamma (IFN-*γ*)^19,20^. Thus, there is a need to adequately describe the dose-responses exhibited by cytokines when administered alone and in combination to establish efficient stimulation regimens.

Despite their ubiquity in a variety of biological contexts, non-monotonic dose-response curves (NMDRCs), also known as biphasic or hormetic dose-response curves, are insufficiently described. In both the literature and in practice, mathematical models are used to quantify dose-response relationships^21-27^. However, conventional approaches fail to accurately describe NMDRCs. Similarly, though numerous mathematical models are used to quantify synergistic and antagonistic drug interactions, they do not correctly capture combinatory effects. To address this shortcoming, Meyer et al.^28^ and Wooten et al.^29^ developed Multi-dimensional Synergy of Combinations (MuSyC), which quantifies synergy based on the law of mass action. This approach distinguishes between synergy types such as potency and efficacy. However, it does not take into consideration NMDRCs.

In this work, we incorporated the principles set forth in MuSyC and developed a new mathematical model, named Uncovering Non-Monotonic Antagonistic and Synergistic Combinations (UNMASC), to describe combinations of agents that exhibit non-monotonic dose-responses. Here, we tested UNMASC by integrating observed dose-response surfaces characterizing *in vitro* T cell progenitor differentiation and maturation following stimulation by combinations of cytokines^9^ (see **Figure 1** for a schematic of our study). We show that UNMASC successfully recapitulates NMDRCs and surfaces, enabling the improved prediction of physiological and drug effects. Additionally, we highlight synergy of potency and efficacy combinations between cytokines that have antagonistic effects alone but that are synergistic in combination, thus providing a rationale for studying these interactions in depth. Thus, UNMASC improves our understanding of cytokine combinations necessary for *in vitro* T cell differentiation and maturation from iPSCs. Thus, this work contributes to establishing efficient stimulation regimens to optimize T cell numbers for CAR-T cell therapy. More broadly, UNMASC can be used to accurately incorporate non-monotonic dose-response surfaces in experimental and clinical contexts, ultimately improving our understanding of the complex phenomena driving such responses.

**Figure 1.**
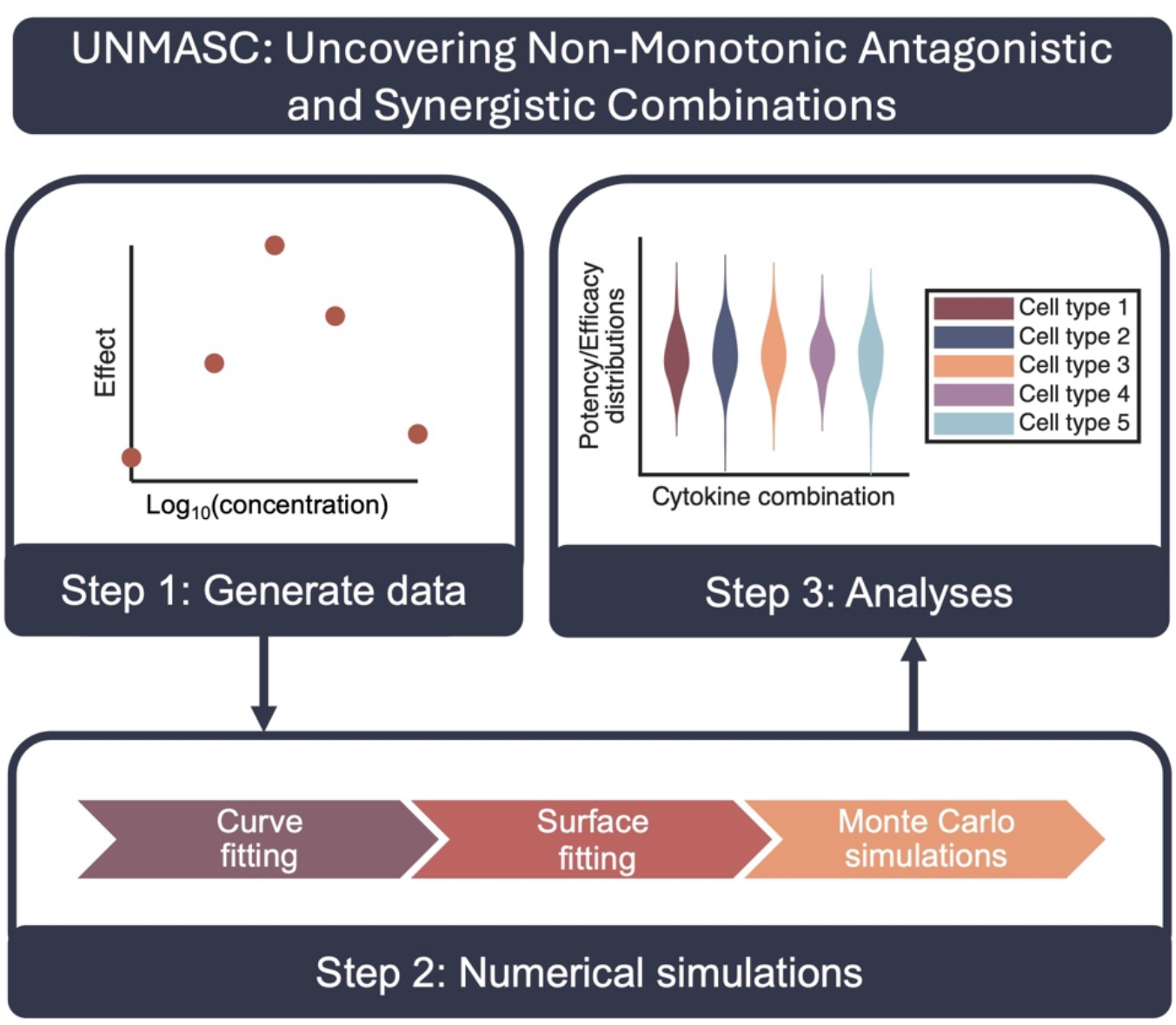
UNMASC: Uncovering Non-Monotonic Antagonistic and Synergistic Combinations. The first step in UNMASC is to experimentally generate data of single and combination dose-response curves and surfaces. In the second step, curve fitting to the dose-responses curves is performed to estimate parameters in the single agent case, before estimating parameters of the dose-response surfaces arising from the combination of multiple agents; parameter distributions are determined by performing Monte Carlo simulations. Lastly, in the third step potency and efficacy parameters are analyzed to determine the effects of the studied combinations on their values.

## METHODS

### Mathematical model describing non-monotonic dose-responses

We generalized MuSyC^28,29^ to account for non-monotonic effects, resulting in a modified Hill equation derived from the Bliss independence principle (see SI). Consider a cell population in which an agent (e.g., cytokine) is introduced. We denote the proportion of cells unaffected and affected by the agent by *U* and *A*, respectively. We are interested in finding the steady state, i.e., when the rate of change of *U* and *A* over time is zero. As laid out in Meyer et al.^28^ and Wooten et al.^29^, the reversible transition between these populations is described by

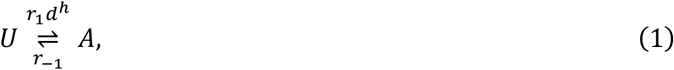

where *d* is the concentration of the agent, *h* is the Hill slope, and *r*_1_ and *r*_−1_ are the forward and backward reaction rate constants, respectively. To account for a biphasic dose-response, let *i* denote the reaction phase (i.e., forward or backward) so that Equation 1 becomes

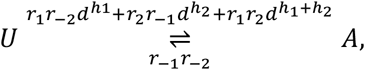

with *h*_*i*_ the Hill slope and *r*_*i*_ the reaction rate for each phase. Mathematically, this biphasic reaction can be written as

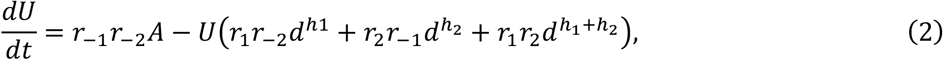

with steady state:

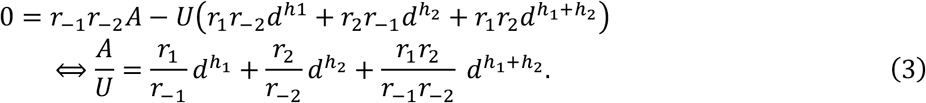

Similar to MuSyC^28,29^, we define 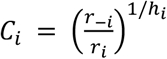 to be the concentration at which half the maximal effect is reached. Therefore, Equation 3 can be rewritten as

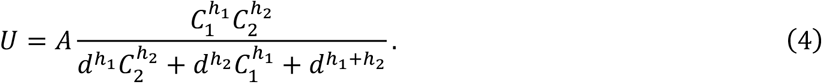

Since cells are either unaffected or affected by the agent, *U* + *A* = 1, Equation 4 can be rewritten as

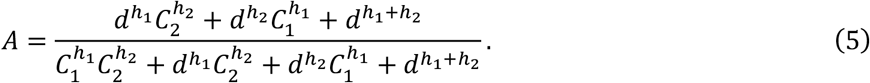

Denoting the baseline effect as *E*_0_ and the effect induced by the agent as *E*_1_, the total effect is given by

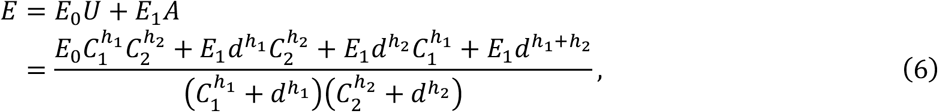

which must satisfy:

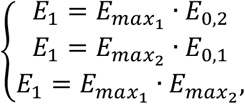

where *E*_0,*i*_ for *i* ∈ {1,2} is the basal efficacy for phase *i* of the dose-response, to reduce to the biphasic Hill equation (see SI). A solution satisfying this system is 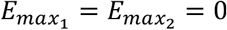, which gives *E*_1_ = 0. Therefore, to describe an increasing then decreasing dose-response, we consider that *E*_1_ is zero by default and that the Hill slope for the first phase is negative.

### Mathematical model describing non-monotonic dose-response surfaces

From the derivation of a single agent’s curve in the previous section, we generalized our method to systems of two agents. We considered cells to be unaffected, affected by agent 1 alone, affected by agent 2 alone, or affected by both agents. The proportion of cells in each of these four states was denoted by *U, A*_1_, *A*_2_ and *A*_1,2_, respectively, giving the system of differential equations:

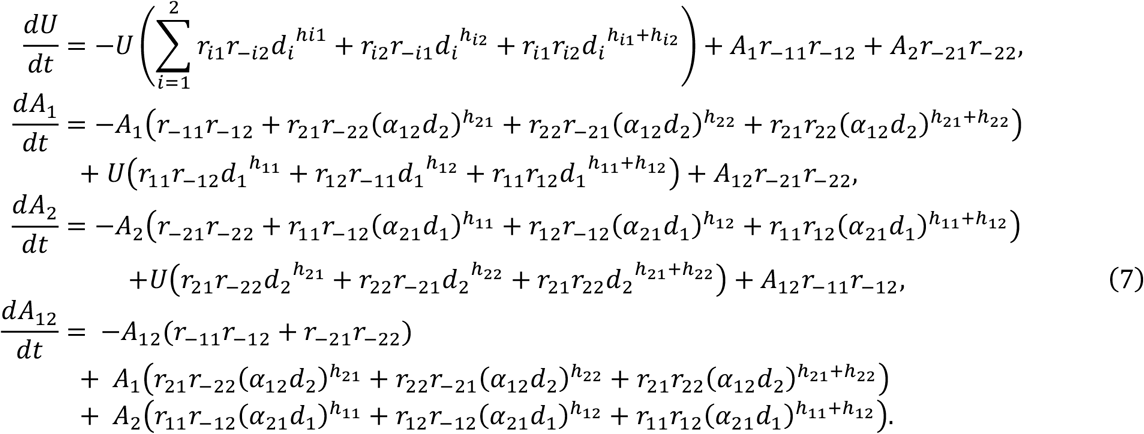

By solving this system at equilibrium and applying the constraint *U* + *A*_1_ + *A*_2_ + *A*_1,2_ = 1 as in MuSyC^28,29^, we obtained

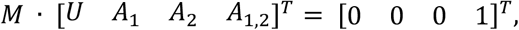

where M is the matrix form of system 7, such that the concentration-dependent effect was found by solving

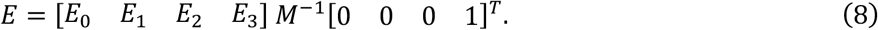

We defined stimulatory relationships as those for which a higher effect is more effective; modifications can be made to account for inhibitory relationships like those observed in e.g., anti-proliferative agents. Note that *E*_0_, *E*_1_, *E*_2_ and *E*_3_ in Equation 8 differ from their meaning in MuSyC^28,29^ because of the imposed negative Hill slope in the first phase of the dose-response of each agent. In our model, the imposed negative Hill slope in the first phase of the dose-response of each agent changed the original meaning of the effect parameters. Accordingly, *E*_0_ denotes the maximal efficacy in the presence of both drugs, with *E*_3_ representing the efficacy in their absence (**Table 1**), and *E*_1_ and *E*_2_ are switched (see **Figure S3**). The framework can describe various surfaces, namely ones in which both agents display increasing and decreasing (I->D) biphasic curves, one agent is biphasic and the other is monotonic, both agents display decreasing and increasing (D->I) biphasic curves, and one agent is I->D and the other is D->I (see **Figure S3**). Thus, to account for the changes in meaning of effect parameters, we defined the synergy of efficacy parameter β to be the increase (or decrease) in efficacy in maximal effect with both agents (*E*_0_ − max(*E*_1_, *E*_2_)) over the most efficacious agent alone (max(*E*_1_, *E*_2_) − *E*_3_) (**Table 1**). As in MuSyC^28,29^, this provided an estimate for the synergy of efficacy. Synergy of potency *α*_*ij*_ was quantified in a similar manner as in MuSyC^28,29^, but adapted to non-monotonic dose-responses (**Table 1**, see **Figure S4** for details).

**Table 1.** Variables and parameters of UNMASC.

| Variable/parameter | Description |
| --- | --- |
| $U$ | Proportion of unaffected cells. |
| $A_i$ | Proportion of cells affected only by agent $i$ ($i \in 1, 2$). |
| $A_{1,2}$ | Proportion of cells affected by both agents. |
| $d_i$ | Concentration of agent $i$ ($i \in 1, 2$). |
| $C_{i,k}$ | Concentration of agent $i$ necessary to achieve 50% of the maximal effect in phase $k$ ($i, k \in 1, 2$). |
| $r_{i,k}, r_{-i,k}$ | Phase $k$ reaction rate of agent $i$ ($i, k \in 1, 2$). |
| $h_{ik}$ | Phase $k$ Hill slope of agent $i$ ($i, k \in 1, 2$). |
| $\alpha_{ij}$ | Fold change in potency of agent $j$ induced by the presence of agent $i$ ($i, j \in 1, 2$). |
| $E_0$ | Maximal combination efficacy. |
| $E_i$ | Maximal efficacy of agent $j$ for $j \neq i$ ($i, j \in 1, 2$). |
| $E_3$ | Efficacy in the absence of treatment. |
| $\beta$ | Increase/decrease in maximal effect with both drugs with respect to the most efficacious single drug ($\beta = \frac{E_0 - \max(E_1, E_2)}{\max(E_1, E_2) - E_3}$). |

### Cytokine dose-responses during T cell development

We previously studied the effects of multiple cytokines, alone and in combination, on *in vitro* T cell progenitor differentiation^9^. In this study, pluripotent stem cells were first differentiated into hemogenic endothelium for 8 days and then taken through the endothelial-to-hematopoietic transition for 7 days (Day -15 to Day 0; see **Figure 3**) . A Central Composite Design (CCD) was used to assess cytokine dose-responses during T cell development (see SI for details). For this, the resulting cells from the hematopoietic progenitor differentiation transition were cultured for 7 days (Day 0 to Day 7; see **Figure 3**) with six cytokines: stem cell factor (SCF), FMS-like tyrosine kinase 3 ligand (Flt3L), interleukin 3 (IL-3), interleukin 7 (IL-7), tumor necrosis factor alpha (TNFa), and stromal-derived factor 1α (CXCL12), and then cultured for fourteen additional days with the same cytokines (Day 7 to Day 21; see **Figure 3**). Absolute cell numbers were measured using flow cytometry after differentiation (days 0-7) and maturation (days 7-21).

**Figure 2.**
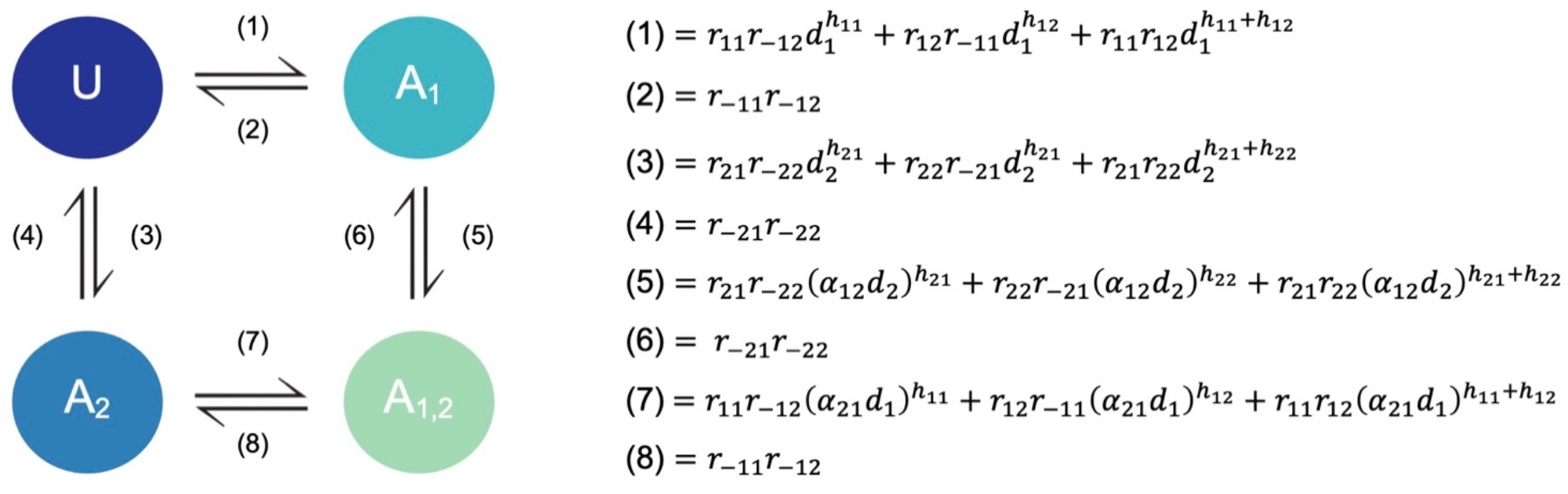
Schematic of the mathematical model used for UNMASC with corresponding forward and backward reaction rates. Conceptualization and reaction rates of unaffected (*U*), affected by agents 1 and 2 (*A*_1_, *A*_2_, respectively), and affected by both agents (*A*_1,2_) from (7).

**Figure 3.**
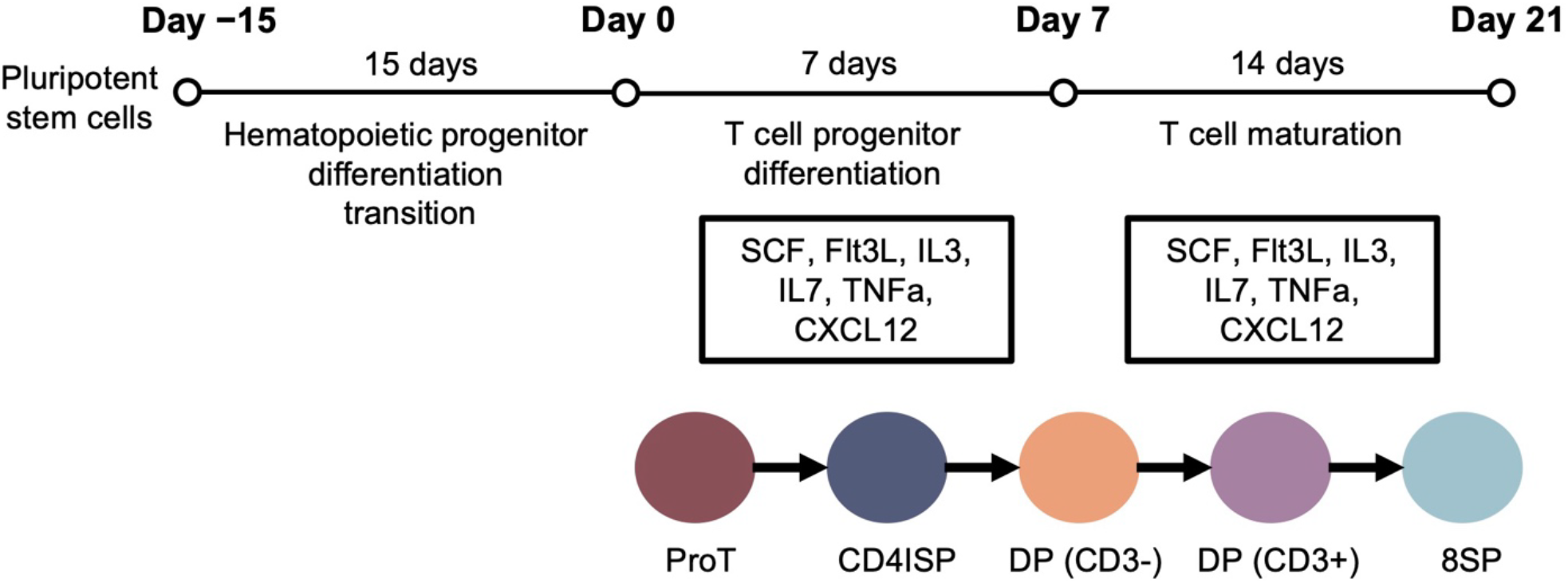
In vitro T cell development assay. Induced pluripotent stem cells (iPSCs) were differentiated into CD34+ hematopoietic progenitors over 15 days. After, cells were cultured for seven days with six cytokines: SCF, Flt3L, IL3, IL7, TNFa and CXCL12. Cells were then tested and cultured again for 14 days with the same cytokines. For the T cell progenitor differentiation, the dose-response curves obtained were for ProT, CD4ISP and DP (CD3-) cells. For the T cell maturation, the dose-response curves obtained were for DP (CD3-), DP(CD3+) and 8SP cells^9^.

For the T cell progenitor differentiation assay (days 0-7), desired cell populations, including CD5+, CD7+ committed T cell progenitors (ProT), CD4+, CD8-, CD3-immature single positive (CD4ISP), and CD4+, CD8+, CD3-double positive (DP (CD3-)) cells were quantified whereas DP (CD3-), DP (CD3+), and CD4-, CD8+, CD3+ single positive (8SP) were measured for the T cell maturation assays from days 7-21 (**Figure 3**). Dose-responses were fit to a polynomial using least squares regression (see SI). The dose-response curve of CXCL12 was not considered during cell maturation (days 7-21) as the polynomial coefficient of degree 1 was not found to be statistically significant (Prob > |t| > 0.05) and CXCL12 was not part of a significant higher order (i.e. quadratic) coefficient^9^. We generated synthetic data points using the polynomial coefficients and the concentrations used for the CCD and normalized them by dividing through the maximal value, so they ranged between 0 and 1.

### Single agent parameter estimation

To estimate individual cytokine effects on differentiation and maturation, we fit the synthetically generated dose-response curves using the in-built Levenberg-Marquardt algorithm in the MATLAB R2024b^30^ *lsqnonlin* function. Each phase of a biphasic curve was fit independently to a typical Hill function (Eq. S1) to ensure each phase had parameters that were described independently from the other phase and the product of the fitted curves provided the biphasic dose-response (Eq. S2). As in MuSyC, a high absolute value of the Hill slope often resulted in numerical errors due to *M* (Eq. 8) being close to singular or badly scaled. Therefore, we limited the range of Hill slopes to [−5,0] or [0,5] by implementing restrictions on the lower and upper bounds of the parameter range, depending on whether the phase’s effect was stimulatory or inhibitory, respectively. We used the polynomial coefficients from the original regression analysis^9^ to calculate data points for each cytokine (see SI for details, with corresponding cytokine concentrations in **Table S1**).

To properly estimate experimental and biological noise, we generated noisy data by sampling from a normal distribution of mean 0 and standard deviation equal to the previously estimated standard errors (SE) of the coefficients in the polynomial fit from Michaels et al^9^. If regression coefficients fell within ± 2 standard errors of the distribution, we kept the coefficients. If at least one of the coefficients fell outside ± 2 standard errors, we discarded both and regenerated new coefficients. We repeated this process to generate a set of 1000 coefficients and then fit all the noisy data to the biphasic or monotonic model (Eq. S1 and S2 in SI) and computed the standard deviation for each parameter distribution. This allowed us to construct confidence intervals about the best-fit parameters^31-33^.

### Estimating combination effects

To estimate the parameters for combination treatments (Eq. 8), we used the concentrations at which half the maximal effect was reached (*C*) and Hill slope estimates from our single agent fits (see previous section) as initial guesses, with bounds defined by the confidence interval from the single agent fits. We then minimized

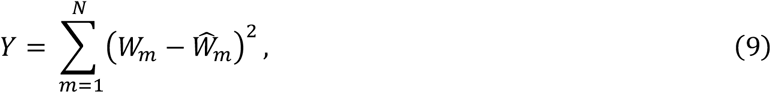

where *N* is the total number of points of the surface, *W*_*m*_ is the *m*^*th*^ data point of the model estimation (Eq. 8) and 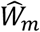 is the *m*^*th*^ data point. *Y* represents the residual norm of the fit, which we normalized by the number of points when assessing the goodness of fit.

## RESULTS

### Non-monotonic dose-responses predicted by UNMASC

Using UNMASC, we first sought to estimate single agent dose-response curves of *in vitro* T cell progenitor differentiation and maturation using 6 cytokines: SCF, Flt3L, IL-3, IL-7, TNFa and CXCL12 (see Methods) to obtain initial estimates for the dose at which half the maximal effect is reached (*C*) and Hill slope parameters of the surface model (Eq. 8). We also set out to generate noise around these dose-response curves that could be used to compute confidence intervals for surface fitting. For this, we generated noise based on our data after CD34+ cells were differentiated into T cell progenitors following stimulation by SCF, Flt3L, IL-3, IL-7, TNFa and CXCL12^9^ (see Methods). In all cases, the monotonic or biphasic behaviour displayed by most noisy trajectories matched with the one present in data points generated using the best-fit coefficients from Michaels et al.^9^ (**Figure S5**). We then fit the dose-response curves for the differentiation stage in days 0-7 and the maturation stage in days 7-21. The number of data points (5) was lower than the number of parameters estimated (8), which could result in a situation where multiple parameter sets could equally well fit the data. However, in practice, models are rarely structurally identifiable and thus rely more on practical identifiability^34^. Our model successfully described the resulting non-monotonic dose-responses, distinguishing inhibitory and stimulatory phases for both differentiation stages (**Figure 4** and **Figure S6**; see **Table S4-Table S9** for estimated parameter values and confidence intervals).

**Figure 4.**
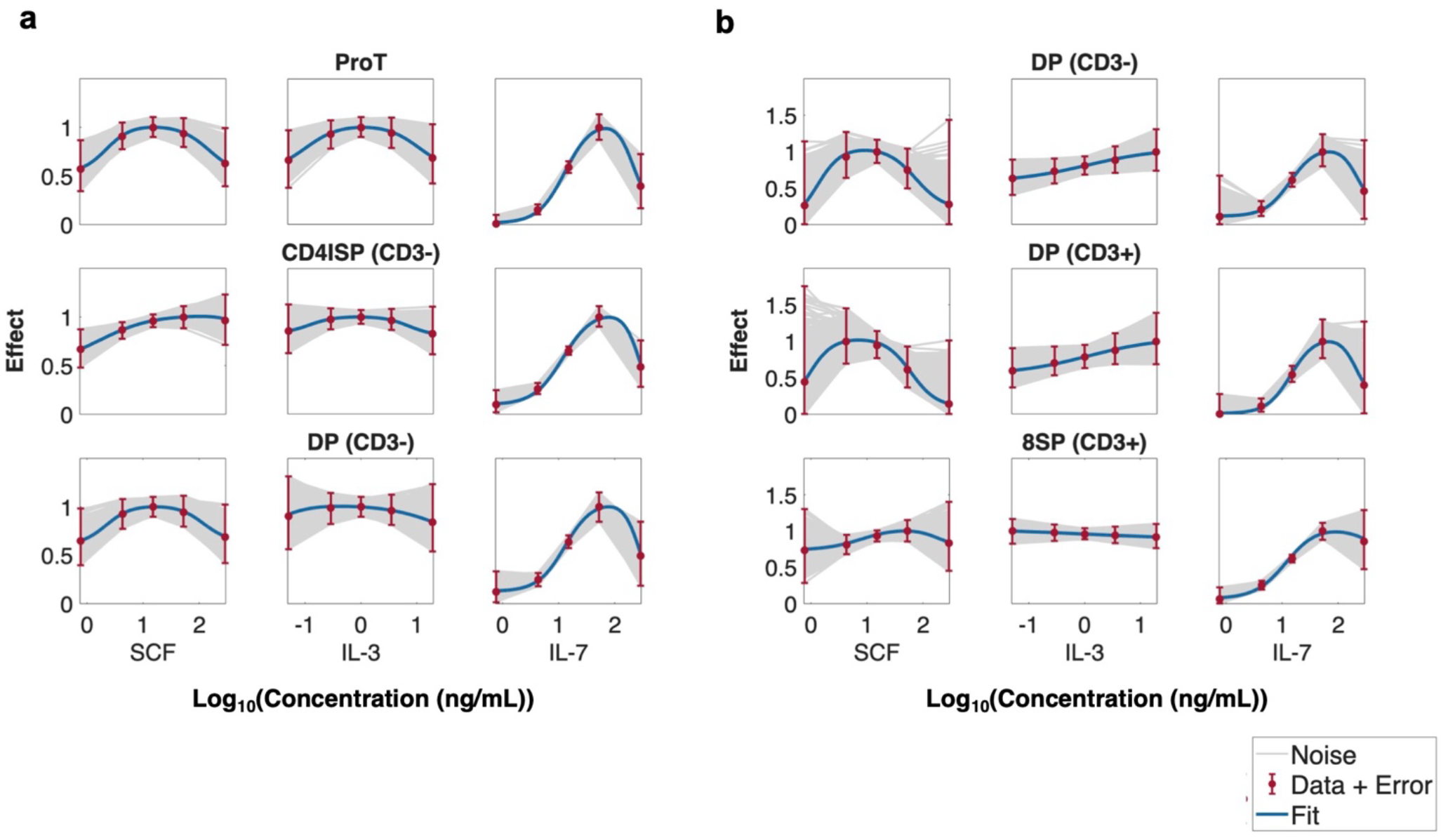
Fits of single agent dose-responses successfully describe non-monotonic behaviours. **a)** Model predictions for cytokines SCF, IL-3 and IL-7 in cell differentiation (days 0-7 of the experiment) for ProT, CD4ISP and DP cells. **b)** Single fits for cytokines SCF, IL-3 and IL-7 in cell maturation (days 7-21 of the experiment) for DP and 8SP cells. **a-b)** Red dots and error bars: measurement ± 2 standard deviations. Blue solid lines: model prediction. Grey lines: generated samples within ± 2 standard deviations of experimental data (see Methods). The measured effect is the cell number normalized by the maximal number of cells.

### UNMASC successfully predicts non-monotonic dose-response combinations

Using the estimates for the dose at which half the maximal effect is reached (*C*) and Hill slope parameters for single agent dose-response curves and the confidence intervals generated, we next sought to assess how accurately UNMASC (Eq. 8) could reproduce the surface dynamics present in the cytokine combinations (see Methods). To do so, we fit our model to our combination cytokine data (**Figure 5** and **Figure S7**). Goodness of fit was defined based on the value of the cost function (Eq. 9 and **Figure S9**) and biological interpretability of the parameters; see **Table S10-Table S15** for all parameter fits. In more than half of the combinations, cytokines exhibited monotonic dose-responses over certain concentration ranges though they were estimated to be monotonic in the single agent fits (**Table S3**). This prevented a good fit in the cell differentiation phase for the TNFa-CXCL12 combination in DP (CD3) cells. Thus, we set TNFa to be monotonic for this interaction, which allowed for an improved estimation (**Figure S7**). The dose at which half the maximal effect was reached (*C*) and Hill slope (*h*) parameter values sometimes varied when estimating dose-response curves alone versus in combinations, but we limited their ranges to the confidence intervals of single fits to ensure they remained within biologically reasonable ranges (see Methods and **Figure S10-Figure S15** for the distributions of noise of single agents and the single and combination fits). Additionally, *α* parameters were set to remain within [10^−4^, 10^4^] and only two combinations had values that went beyond 10^3^ (TNFa-CXCL12 in CD4ISP cells and IL7-TNFa in DP (CD3-) cells, both in cell differentiation (days 0-7) (see **Table S11** and **Table S12**)). Overall, our model successfully captured the biphasic behaviours present in the cytokine combinations. In particular, effect parameters (*E*_*i*_) were successfully estimated to represent the maximal/minimal effect (**Figure 5; Figure S7** and **Table S10-Table S15**).

**Figure 5.**
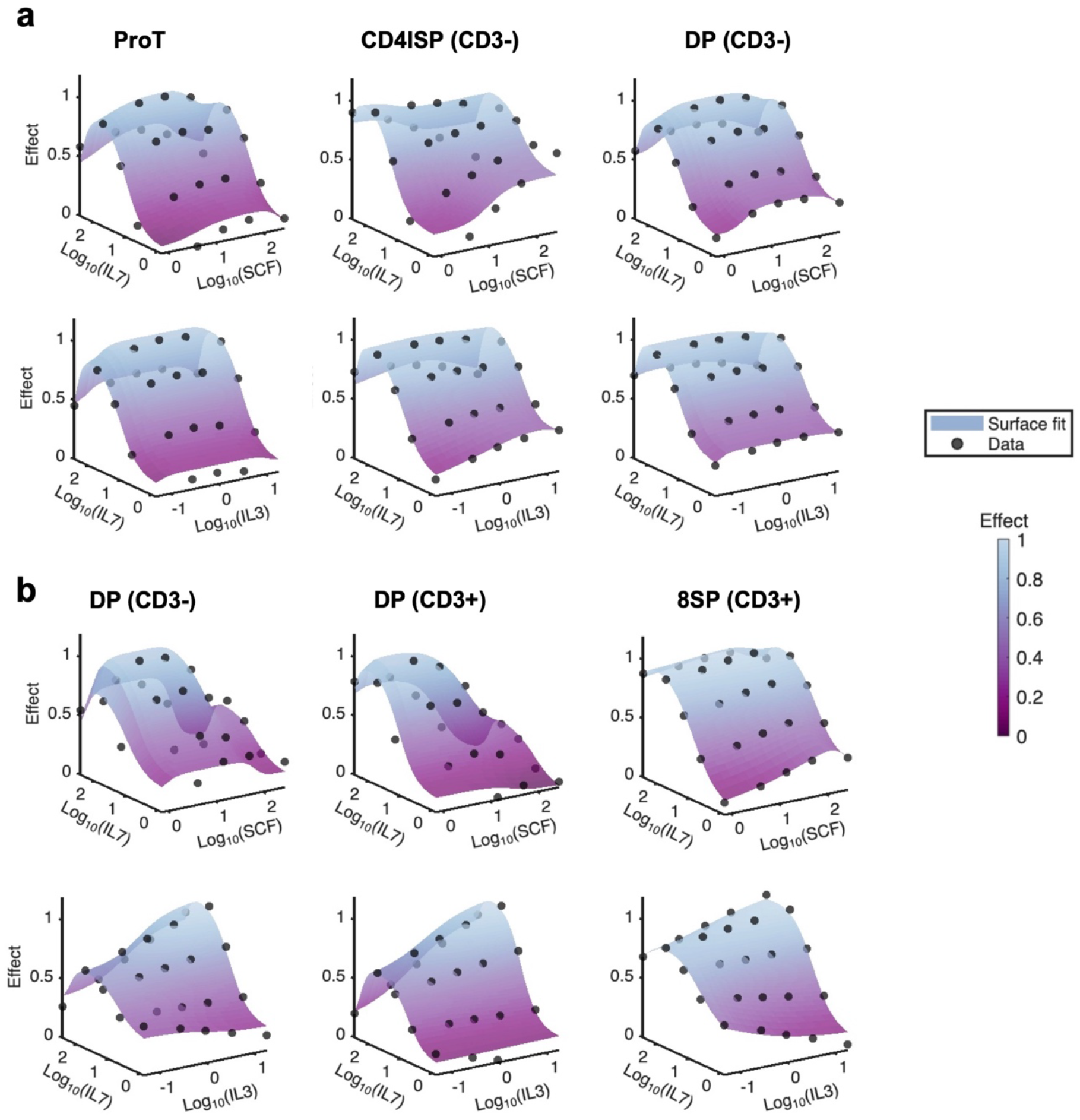
Combination surface fits successfully recapitulate non-monotonic behaviours. **a)** Model predictions of cytokine combinations SCF-IL7, IL3-IL7, IL7-TNFa and TNFa-CXCL12 in cell differentiation (days 0-7 of the experiment) for ProT, CD4ISP and DP cells. **b)** Model predictions of cytokine combinations SCF-IL7, IL3-IL7 and IL7-TNFa in cell maturation (days 7-21 of the experiment) for DP and 8SP cells. **a-b)** Black dots: data generated from polynomial regression coefficients. Scaled purple surfaces: combination fits with our new model. The axes for the cytokine concentrations are the physical concentrations from the previous experiment scaled with a logarithm of base 10 (**Table S1**). The vertical axis is the effect (between 0 and 1).

### UNMASC suggests synergistic potency of IL-7 during differentiation due to the presence of SCF and IL-3

We then investigated how potency and efficacy changed in combination versus as single agent. This is critical for determining target combinations of interest for continued development. For this, we ran 200 Monte-Carlo simulations by sampling from a normal distribution centered at 0 and standard deviation equal to the square root of the normalized residual norm of the best fit at each data point, without limiting their range. We then fit to each resulting surface fits using best-fit parameters as initial guesses. This procedure allowed us to define confidence intervals to estimate the distributions of *α*_*ij*_ and *β*.

During differentiation, we found that IL-7 acts as a potency antagonist for SCF and IL-3. Indeed, the mean log fold change of potency with standard deviation of IL-7 on SCF was -0.56 (0.07) for ProT cells, -0.69 (0.13) for CD4ISP cells and -0.53 (0.03) for DP (CD3-) cells (**Table S16** and **Figure 6a**). In the same period, the mean log fold change of potency of IL-7 on IL-3 was -0.55 (0.07) for ProT cells, -0.36 (0.06) for CD4ISP cells and -0.73 (0.25) for DP (CD3-) cells (**Table S16** and **Figure 6a**). Interestingly, we found that SCF acts as a potency agonist for IL-7 during differentiation in ProT and DP (CD3-) cells, with a mean log fold change of potency of 0.72 (0.85) and 0.36 (0.06) respectively (**Table S16** and **Figure 6b**). Similarly, IL-3 was determined to act as a potency agonist for IL-7 in ProT, CD4ISP and DP (CD3-) cells during differentiation, with a mean log fold change of potency of 1.37 (1.46), 0.06 (0.73) and 1.30 (0.88) respectively (**Table S16** and **Figure 6b**). We also found that TNFa acts as a potency agonist for IL-7 in ProT, CD4ISP and DP (CD3-) cells (**Table S16** and **Figure S16a**).

**Figure 6.**
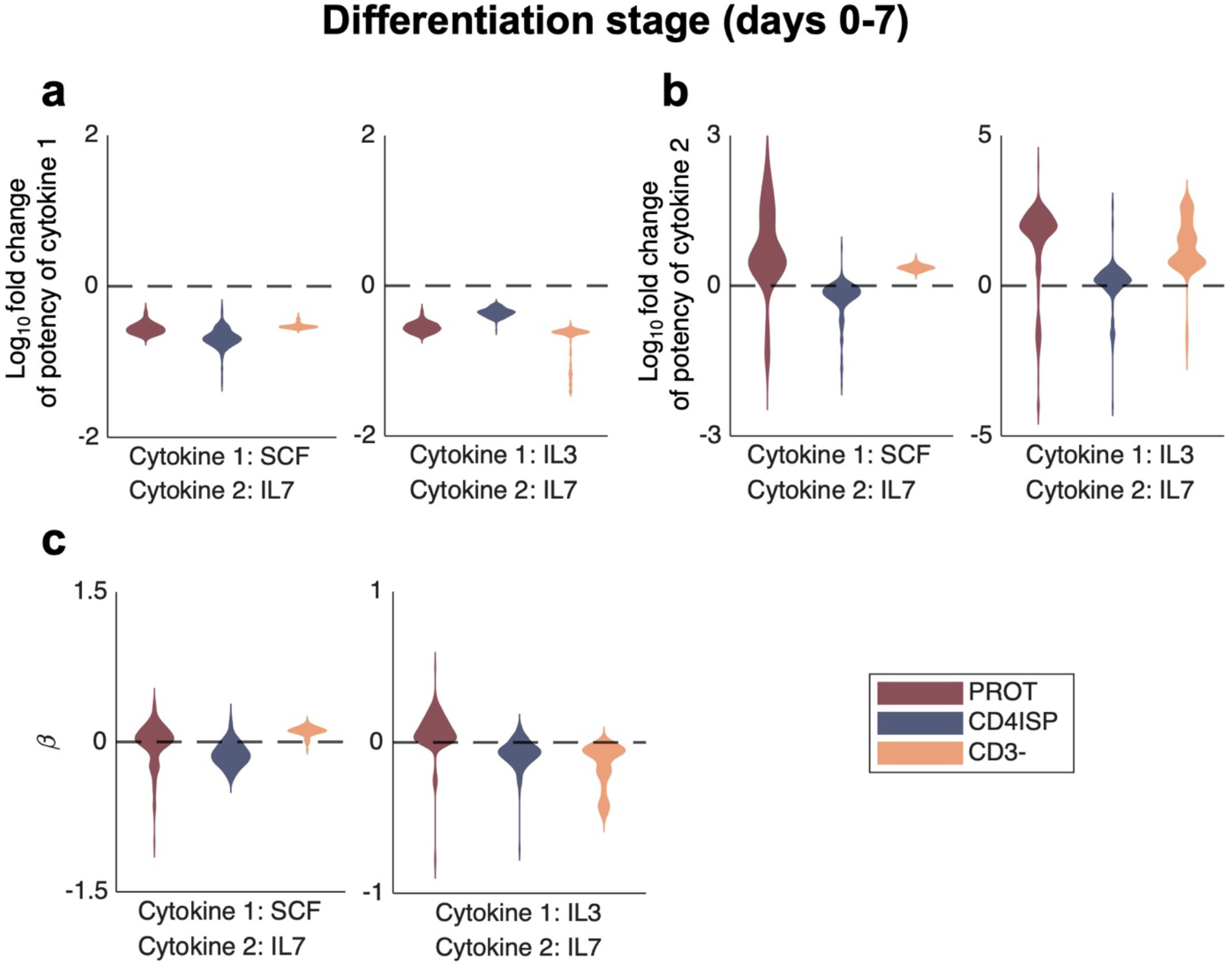
Distributions of potency and efficacy parameters in cytokine combinations during differentiation (days 0-7) following Monte Carlo simulations. **a)** Distributions of the log_10_ fold change in potency of the first cytokine due to the presence of the second cytokine for combinations SCF-IL7 and IL3-IL7 in days 0-7 of the experiment. **b)** Distributions of the fold change in potency of the second cytokine due to the presence of the first cytokine for combinations SCF-IL7 and IL3-IL7 in days 0-7 of the experiment. **c)** Distributions of synergy of efficacy parameters (β) for cytokine combinations SCF-IL7 and IL3-IL7 in days 0-7 of the experiment. **a-c)** First and second cytokines for each combination are shown on the x-axis (first cytokine on top and second cytokine below). Dotted line: no synergy of potency/efficacy. Above and below the dotted line denote synergistic and antagonistic potency/efficacy respectively.

To assess how different cytokine combinations affect the synergy of efficacy during differentiation (days 0-7), we analyzed the distributions of *β*. We found little to no synergy of efficacy for combinations involving SCF-IL7 and IL3-IL7 (**Table S16** and **Figure 6a**). This is likely due to the already high effect of IL7 alone. During the same period, the TNFa-CXCL12 interaction was found to show antagonistic synergy of efficacy for all cells during the differentiation phase with means of -1.52 (0.31) for ProT cells, -1.76 (0.18) for CD4ISP cells and -3.53 (0.34) for DP (CD3-) cells (**Table S16** and **Figure S16a**).

### UNMASC suggests synergy of efficacy of IL3-IL7 combination during maturation

During maturation (days 7-21), we again found that IL-7 acts as a potency antagonist for SCF in DP (CD3- and CD3+) cells (mean log fold change of -0.62 (0.16) and -0.41 (0.77) respectively), (**Table S17** and **Figure 7a**)) and as a potency antagonist for IL-3 in DP (CD3- and CD3+) and 8SP cells (mean log fold change of 0.85 (0.06), -0.73 (0.29) and 1.03 (0.53) respectively, (**Table S17** and **Figure 7a**)). The positive log fold change in the potency in IL-3 is indicative of an antagonistic potency, since IL-3 decreased DP (CD3-) and 8SP cells (**Table S3**). Next, we again observed that SCF and IL-3 act as synergy agonists on IL-7 in DP (CD3- and CD3+) and 8SP cells (see **Table S17** and **Figure 7b**). Further, we also found that TNFa acts as a potency agonist for IL-7 (**Table S17** and **Figure S17A**) and that IL-7 acts as a potency antagonist for TNFa (**Table S17** and **Figure S17b**). Together, these results further suggest that IL-7 is a potency antagonist for SCF, IL-3 and TNFa, while SCF, IL-3 and TNFa are potency agonists for IL-7. We also observed synergistic efficacy between IL-3 and IL-7 (**Figure 7c**) across all cell types during the maturation phase, with means of 1.19 (0.33) for DP (CD3-), 1.51 (0.56) for DP (CD3+) and 0.36 (0.20) for 8SP cells (**Table S17**), despite IL-3 being monotonic decreasing in DP (CD3-) and SP (CD3+) cells (**Figure 5b** and **Figure S8**).

**Figure 7.**
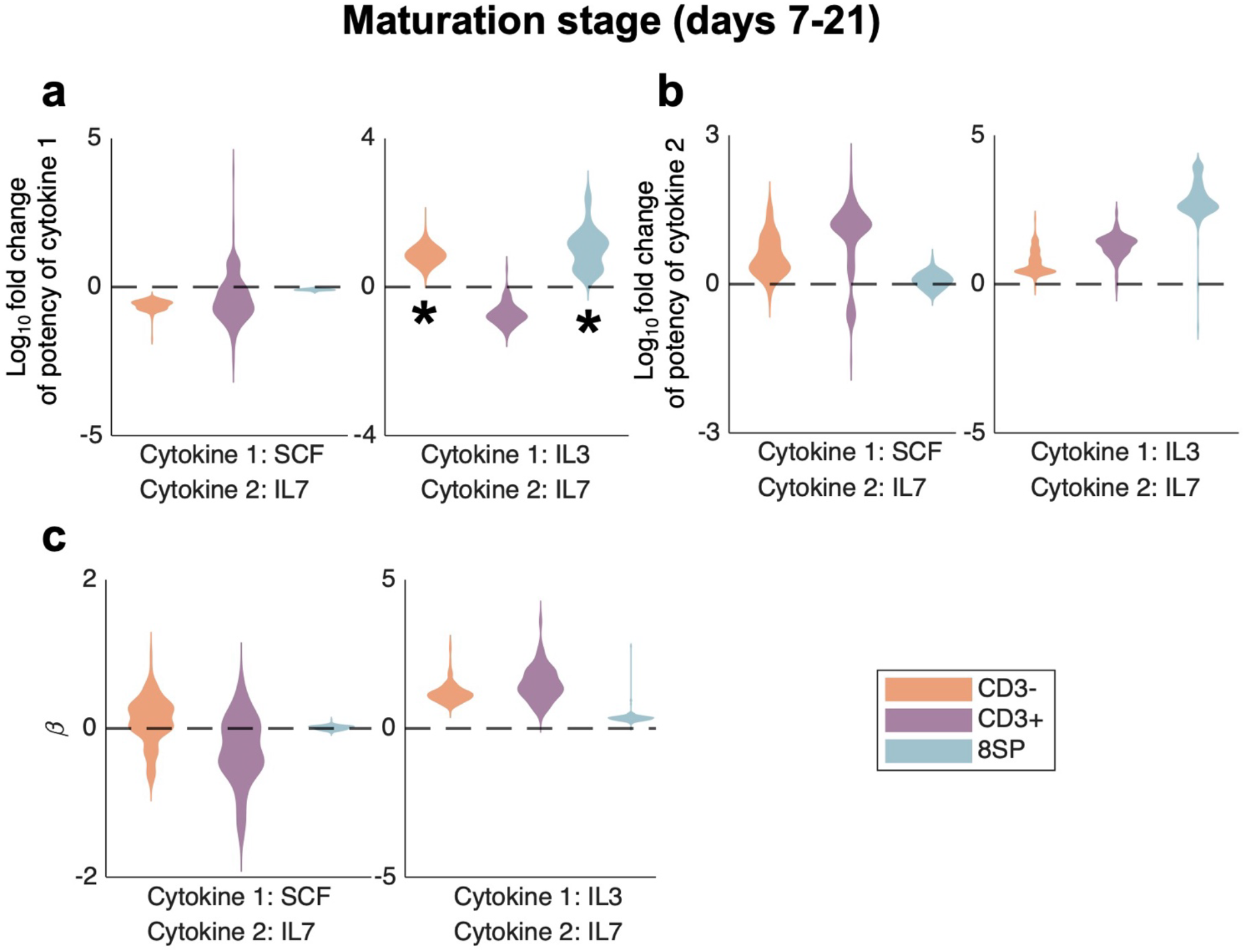
Distributions of potency and efficacy parameters in cytokine combinations during maturation (days 7-21) following Monte Carlo simulations. **a)** Distributions of the fold change in potency of the first cytokine due to the presence of the second cytokine for combinations SCF-IL7 and IL3-IL7 in days 7-21 of the experiment. **b)** Distributions of the fold change in potency of the second cytokine due to the presence of the first cytokine for combinations SCF-IL7 and IL3-IL7 in days 7-21 of the experiment. **c)** Distributions of synergy of efficacy parameters (*β*) for cytokine combinations SCF-IL7 and IL3-IL7 in days 7-21 of the experiment. **a-c)** First and second cytokines for each combination are shown on the x-axis (first cytokine on top and second cytokine below). Dotted line: no synergy of potency/efficacy. Above and below the dotted line denote synergistic and antagonistic potency/efficacy respectively. Black asterisk: distributions in which above the dotted line denotes antagonistic potency and below zero denotes synergistic potency due to the decreasing behaviour of the curve.

## DISCUSSION

In this work, we developed UNMASC, a new method to fit dose-response surfaces for combinations of agents displaying non-monotonic behaviours. UNMASC estimates parameters with direct biological interpretations and distinguishes between synergistic efficacy and synergistic potency. We tested UNMASC on experimental dose-response surfaces that displayed non-monotonic behaviour and successfully captured the nonmonotonicity of observed surfaces and distinguished between synergy types.

Our analysis of *in vitro* dose-response surfaces suggests that IL-7 interacts with SCF, IL-3 and TNFa with synergistic potency. Additionally, during maturation, IL3-IL7 was found to have synergistic efficacy, despite the fact that IL-3 displayed monotonic decreasing dose-responses for certain cell types. This underlines the importance of incorporating agents that display decreasing dose-responses in combination experiments, as their combined effect can be more efficacious than that of the one of agents alone. Additionally, the difference in monotonic behaviour (i.e., increasing or decreasing) for IL-3 when alone or in combination highlights the need to test combinations of agents across a wide range of concentrations to assess when changes in dose-response behaviours occur.

Since UNMASC only has one basal efficacy parameter and does not consider monotonicity directional changes, estimated dose response surfaces will always go back to their basal efficacy at high concentrations (**Figure S8**). This can be a limitation when, for example, a substance at high dosage is more inhibitory than the baseline effect. Additionally, in more than half of combinations, substances that displayed a biphasic behaviour alone exhibited a monotonic behaviour when in combination, sometimes only for certain concentrations (**Table S3**). This could be due to several factors, namely if the initial hypothesis that the substance alone displayed a biphasic dose-response was erroneous due to noisy data or lack of sufficient data for the interaction surface to accurately describe the mechanism. In these cases, solving the system in Eq. 8 can result in numerical instability. However, solving the system while assuming the data are monotonic still allows an analysis of the surface, like in the TNFa-CXCL12 interaction in DP (CD3-) cells during differentiation in days 0-7. Further, the noise incorporated to experimental data was estimated to be independent and symmetric at each measure^31,33^, which is often not the case in practice^35,36^.

Though existing, simpler methods like polynomial and spline fits can be used to characterize surfaces of interaction for substances that display non-monotonic behaviours, such methods cannot provide biological meaning for underlying mechanisms driving these responses. UNMASC yields a clear rationale for estimated parameters as they are directly linked to the monotonic Hill equation. Further, the estimated noise provides a quantitative measure of the uncertainty in the parameters. While this noise does not account for the domain of the data, the bounds of fitted parameters were restricted, thus ensuring they were estimated to realistic values. By decoupling synergistic potency and efficacy, as in MuSyC^28,29^, we can obtain more information on how cytokines interact *in vitro*, with generalization to other substances. Further, our model can easily allow for the differentiation of these two types of synergy when considering surface interactions that display non-monotonic behaviours. Thus, our new framework to estimate non-monotonic combination dose-responses can in turn be used for informed decision-making when designing studies and optimizing the doses. This work therefore represents a significant step-forward in the study of non-monotonic dose-response interactions with applications to a wide variety of biological contexts.

## Data availability

The data analyzed in this study are available from Michaels et al. 10.1126/sciadv.abn5522, Tables S1-S7.

## Code availability

All code required for recreating manuscript analyses are available for review at https://github.com/Craig-Lab/UNMASC.git.

## Contributions

GBG, YM, and MC conceived and designed the research. GBG and MC performed the research. GBG analyzed the results. GBG, YM, and MC wrote and edited the manuscript.

