## Supplementary Information for "Uncovering Non-Monotonic Antagonistic and Synergistic Combinations (UNMASC), a robust method with applications to T cell differentiation *in vitro*"

#### 1. Non-monotonic dose-response curves

The typical Hill equation describing monotonic dose-responses can be written as

$$E = E_0 + \frac{E_1 - E_0}{1 + (C/d)^h}, \tag{S1}$$

where $E_0$ and $E_1$ correspond to the effect in the absence of the drug and maximal effect of the drug, respectively, $C$ denotes the concentration at which half the maximal effect is obtained, and $h$ is the Hill slope (**Figure S1a**). Various models[1-4] have attempted to model non-monotonic dose responses based on Eq. $S1$. However, most do not provide biological meaning for all model parameters. Conversely, Beckon et al. developed a biologically-motivated model by considering the positive and negative effects of the biphasic curve to have distinct sensitivity threshold probability distributions[5]. In this vein, consider the modification of the Hill equation above by describing each phase as an independent process (**Figure S1b**). Then the entire dose-response can be written as the product of these features[6]. In the case of a biphasic dose-response, this gives

$$E = \left(E_{0,1} + \frac{E_{max_1} - E_{0,1}}{1 + (C_1/d)^{h_1}}\right)\left(E_{0,2} + \frac{E_{max_2} - E_{0,2}}{1 + (C_2/d)^{h_2}}\right). \tag{S2}$$

This equation is based on the Bliss Independence model which assumes that the resulting effect is the multiplication of the effects[7]. It is important to note that the intersection of the two independent processes will impact the shape of the resulting effect curve. Since Equation $S2$ only holds meaning if the data is normalized, $E_{max_i} = 1, i \in \{1,2\}$. Having two curves that do not intersect at the peak can lead to maximal effects that do not reach 1 and ultimately have an impact on the maximal effect of the dose-response (**Figure S2**).

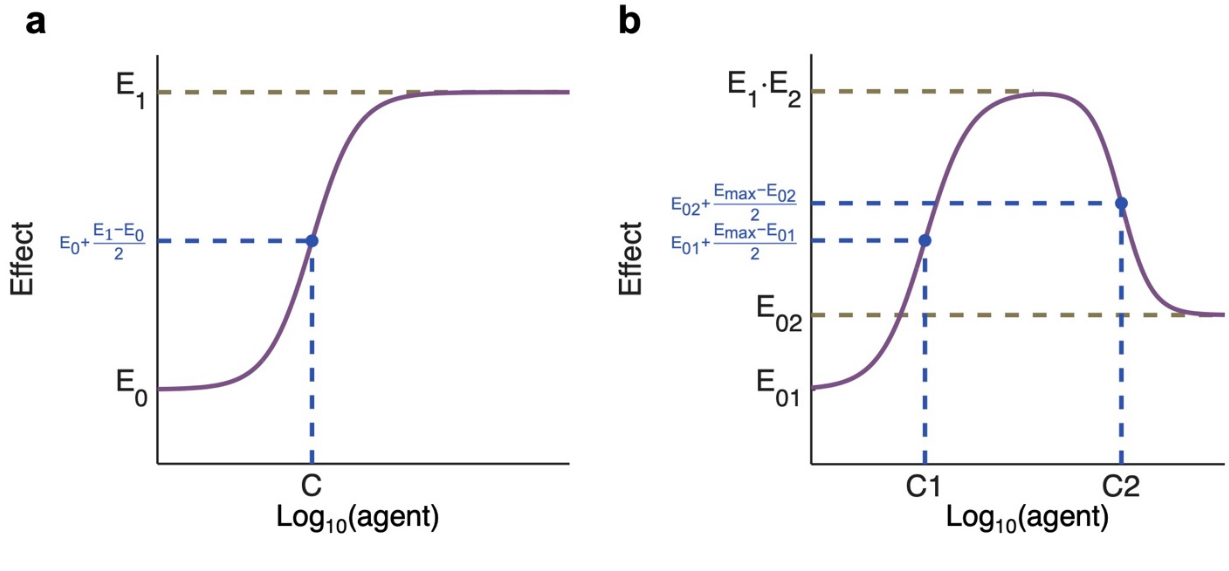


**Figure S1. Monotonic and biphasic dose-response curves based on the Hill equation. a)** Stimulatory (increasing) saturable dose-response described by the typical Hill equation (Eq. S1). $E_1$ maximal effect. $E_0$: baseline effect. C: dose at which half the maximal effect is reached. **b)** Biphasic dose-response with the same parameters as a typical Hill curve (Eq. S1), as described by Eq. S2.

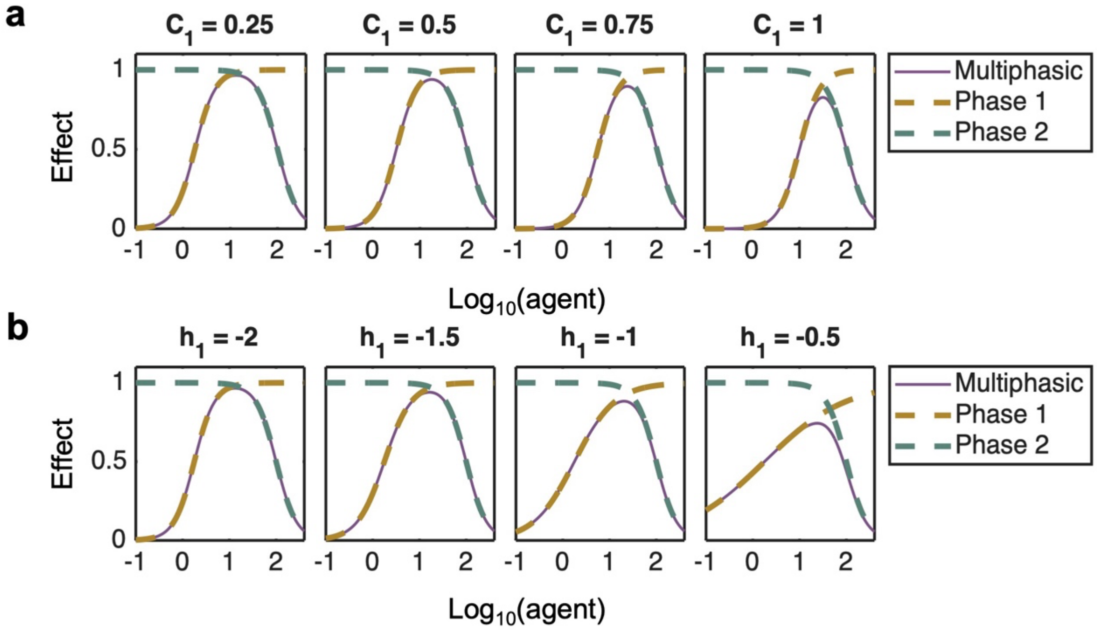


**Figure S2. Impact of parameters of the biphasic dose-response model. a)** Biphasic dose-response curves that vary in their EC50 values for the first phase (i.e. parameter $C_1$). Other parameters are fixed: $E_{01} = E_{02} = 1$, $E_1 = E_2 = 0$, $h_1 = -2$, $h_2 = 2$, $C_2 = 2$. **b)** Biphasic dose-response curves that vary in the Hill slope for the first phase. Other parameters are fixed: $E_{01} = E_{02} = 1$, $E_1 = E_2 = 0$, $h_2 = 2$, $C_1 = 0.25$, $C_2 = 2$.

### 2. Non-monotonic dose-response surfaces

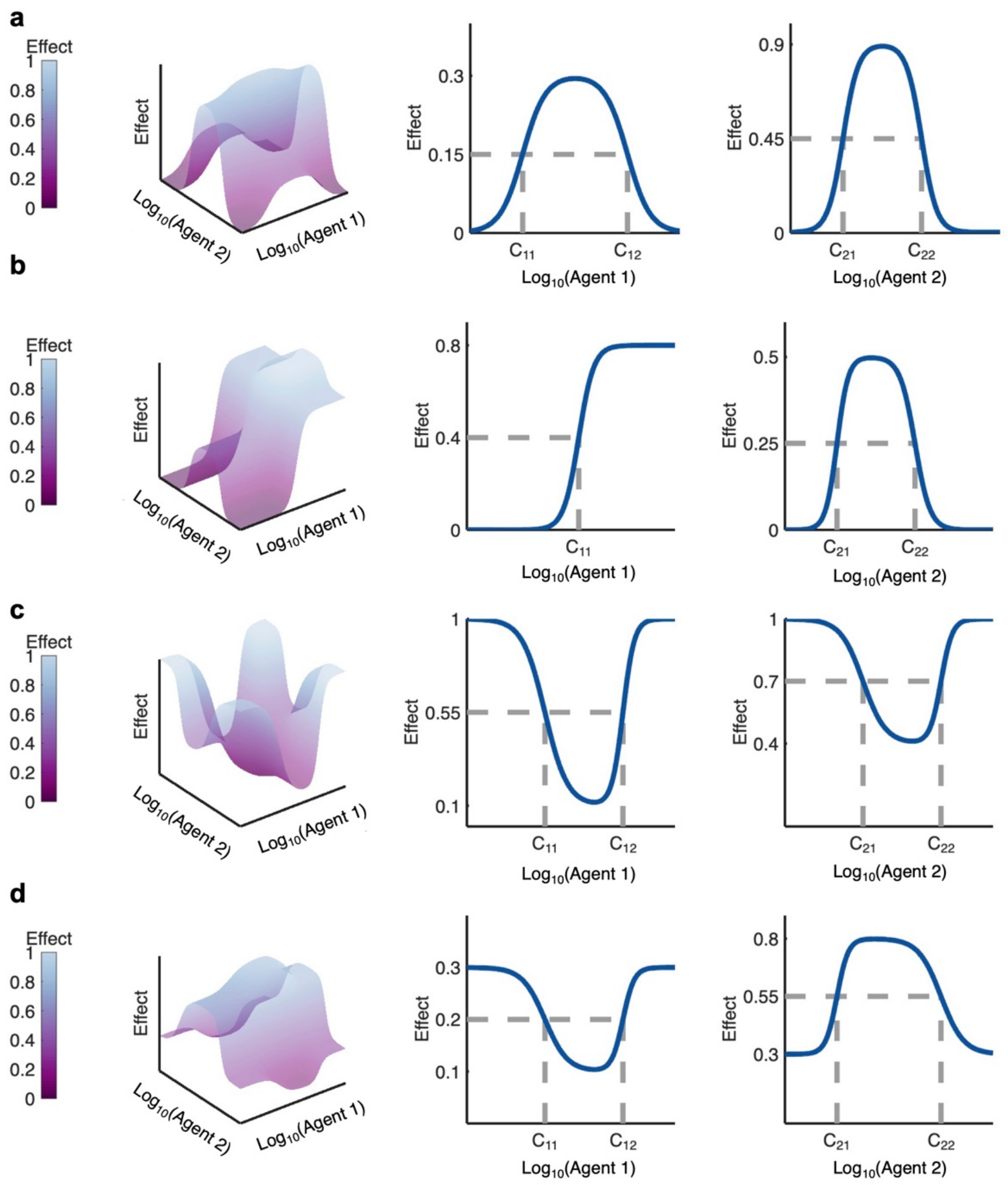


**Figure S3. Behaviours of different surface types in the new model.** Left: combination effect surface. Middle: dose-response of agent 1 when $d_2 \to 0$. Right: dose-response of agent 2 when $d_1 \to 0$. **a)** Interaction between two agents that display increasing-decreasing (I->D) dose-responses. Parameters values used: $E_0 = 1$, $E_1 = 0.9$, $E_2 = 0.3$, and $E_3 = 0$. **b)** Interaction between an agent that displays monotonic behaviour and an agent that displays biphasic I->D behaviour. Parameters values used: $E_0 = 1$, $E_1 = 0.5$, $E_2 = 0.8$, and $E_3 = 0$. **c)** Interaction between two agents that display decreasing-increasing (D->I) dose-responses. Parameters values used: $E_0 = 0$, $E_1 = 0.4$, $E_2 = 0.1$, and $E_3 = 1$. **d)** Interaction between an agent that displays biphasic D->I behaviour and an agent that displays biphasic I->D behaviour. Parameters values used: $E_0 = 1$, $E_1 = 0.8$, $E_2 = 0.1$, and $E_3 = 3$.

### 3. Synergy and antagonism of potency in non-monotonic dose-responses

To assess synergistic and antagonistic potency for non-monotonic combinations, we redefined the concept introduced in MuSyC for monotonic dose-responses[8,9]. By definition, non-monotonic combination relationships exhibit increasing and decreasing phases. In our model, parameter $\alpha_{ij}$ quantifies the fold change in potency of agent $j$ induced by agent $i$ (i.e., the change in concentration at which the half-maximal effect of one agent is reached ($C$) due to the presence of another). Dose-responses that are biphasic have two $C$ values, but we considered the same $\alpha$ parameter for both. Depending on the value of $\alpha_{ij}$, this results in a shift to the right or to the left of the entire dose-response. During a dose-response that is either monotonic increasing or an increasing then decreasing biphasic response (I→D), $\alpha_{ij} < 1$ represents antagonist potency and $\alpha_{ij} > 1$ denotes synergistic potency (**Figure S4a**). However, in a monotonic decreasing or biphasic decreasing then increasing dose-response (D→I), $\alpha_{ij} < 1$ instead denotes a higher $C$, and thus a higher concentration is required before the effect eventually decreases. Since we consider a higher effect to be more effective, we consider $\alpha_{ij} < 1$ in this case to be synergistic potency. Similarly, $\alpha_{ij} > 1$ indicates a smaller $C$ and therefore a lower concentration before the effect decreases, which we define as antagonistic potency (**Figure S4b**).

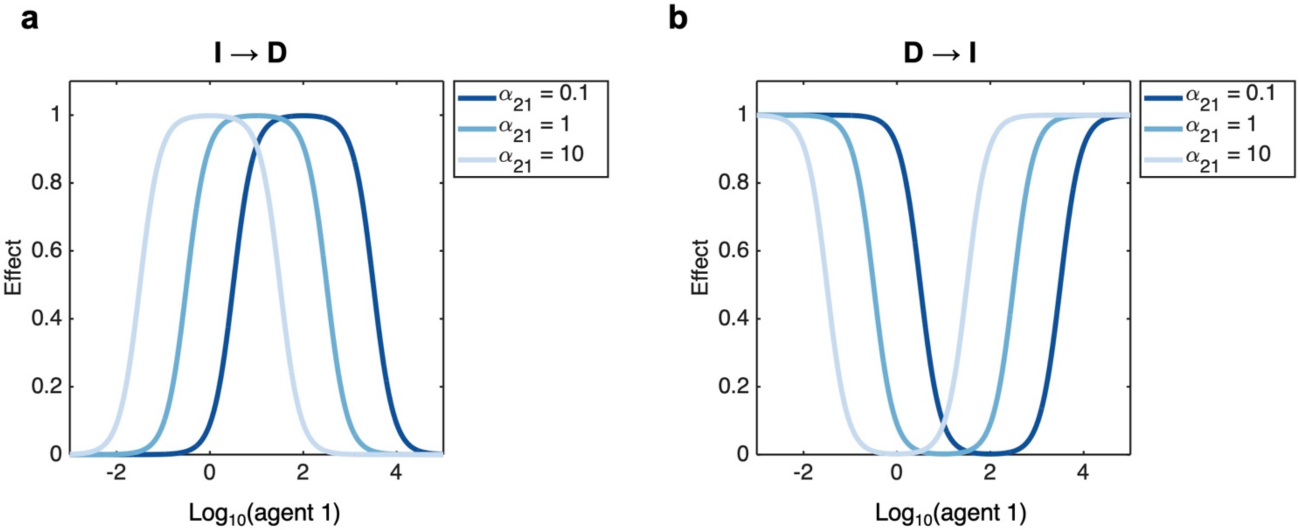


**Figure S4. Interpretation of synergistic potency depends on increasing/decreasing behaviour of the dose-response curve. A)** Effect of parameter $\alpha_{21}$(fold change in the potency of agent 1 induced by the presence of agent 2) on the effect of agent 1 in the case of an I->D dose-response. $\alpha = 0.1$ represents antagonistic potency, $\alpha = 1$ represents no change in potency and $\alpha = 10$ represents synergistic potency. **B)** Effect of parameter $\alpha_{21}$ (fold change in the potency of agent 1 induced by the presence of agent 2) on the effect of agent 1 in the case of a D->I dose-response. The D->I dose-response is obtained by having 1-E, where E is the dose-response of an I->D curve. $\alpha = 0.1$ represents synergistic potency, $\alpha = 1$ represents no change in potency and $\alpha = 10$ represents antagonistic potency.

### 4. Data preparation

In Michaels et al.[10] a Central Composite Design (CCD) was used to assess cytokine dose-responses during T cell development. Pluripotent stem cells were differentiated through hematopoietic progenitor differentiation for 15 days (days -15 to 0). Resulting cells were cultured for seven days (days 0-7) with stem cell factor (SCF), FMS-like tyrosine kinase 3 ligand (Flt3L), interleukin 3 (IL-3), interleukin 7 (IL-7), tumor necrosis factor alpha (TNFa), and stromal-derived factor 1α (CXCL12) before being cultured for fourteen additional days (days 7-21) with the same cytokines. Absolute cell numbers were measured using flow cytometry and the dose-response model:

$$Y = \beta_0 + \sum_{i=1}^{k} \beta_i X_i + \sum_{i=1}^{k-1} \sum_{j=i+1}^{k} \beta_{ij} X_i X_j + \sum_{i=1}^{k} \beta_{ii} X_{ii}^2 + \varepsilon, \quad (S3)$$

was fit for the cell differentiation phase (days 0-7) and the cell maturation phase (days 7-21) using least-squares regression. In the equation above (Eq. $S3$) , $Y$ represents the square root of the number of cells, $X_i$ the concentration of cytokine $i$, $\beta_i$ denotes the linear coefficient of cytokine $i$, $\beta_{ii}$ the quadratic coefficient of cytokine $i$, and $\beta_{ij}$ for $i \neq j$ the interaction term between cytokine $i$ and cytokine $j$. Scaled levels of concentrations for the CCD were given by

$$x = [-2.366, \ -1, \ 0, \ 1, \ 2.366].$$

The conversion to physical concentrations in ng/mL was obtained using the following transformation

$$c = c_0 \cdot 3.5^x,$$

where $c_0$ is the concentration when $x = 0$. The value of $c_0$ for each cytokine was set to ensure that concentrations are within a reasonable range. Finally, the exponent base was selected to ensure a sufficiently wide range of cytokine concentrations so that the change in response would be greater than measurement noise or biological variation[10,11].

Our model requires normalizing the data so that they range between 0 and 1. For this, we generated data points from the previously estimated polynomial fits using the reported regression coefficients. The number of cells was obtained by squaring this value. We normalized the resulting number of cells by dividing by its maximal value over the concentration range. We then transformed the scaled concentrations to physical concentrations (in $\log_{10}$ concentrations) to perform single agent fits. When generating data points for the surfaces, we rejected values that were negative, only keeping values that were positive or null. Of the 21 surfaces analyzed, 9 contained at least one negative data point, with no more than 4 negative data points observed in any of these cases.

| | | **CONCENTRATION (ng/mL)** | | | | |
|---|---|---|---|---|---|---|
| **CYTOKINE** | SCF | 0.77 | 4.29 | 15 | 52.5 | 290.64 |
| | Flt3L | 0.52 | 2.86 | 10 | 35 | 193.76 |
| | IL3 | 0.05 | 0.29 | 1 | 3.5 | 19.38 |
| | IL7 | 0.77 | 4.29 | 15 | 52.5 | 290.64 |
| | TNFa | 0.02 | 0.11 | 0.4 | 1.4 | 7.75 |
| | CXCL12 | 0.77 | 4.29 | 15 | 52.5 | 290.64 |

**Table S1. Cytokine concentrations from Michaels et al.** Data points generated from the polynomial coefficients were evaluated at each of these concentrations to obtain all data points in single dose-responses and surfaces[10].

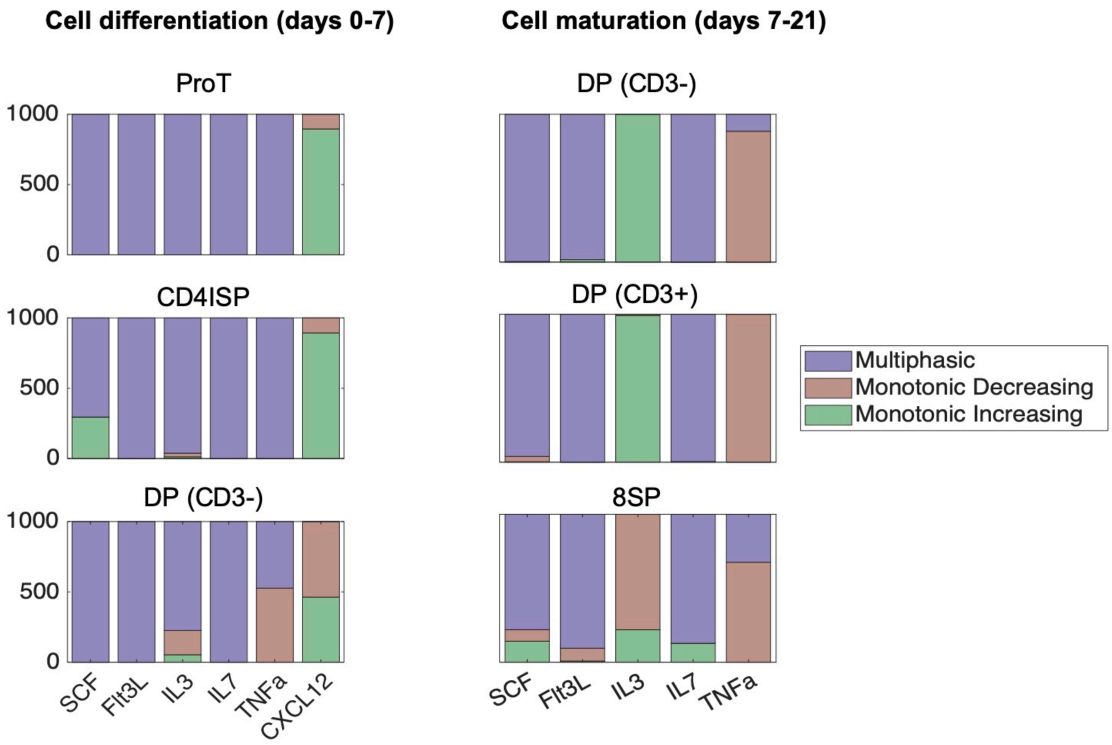


**Figure S5. Noise contribution for single dose-responses.** Noise was classified based on behaviour (multiphasic, monotonic decreasing, or monotonic increasing), the cytokine, and the cell type. For surface fits, we selected the behaviour most frequently observed in the generated data for the corresponding cytokine and cell type over time.

### 5. Cytokine combination and fits

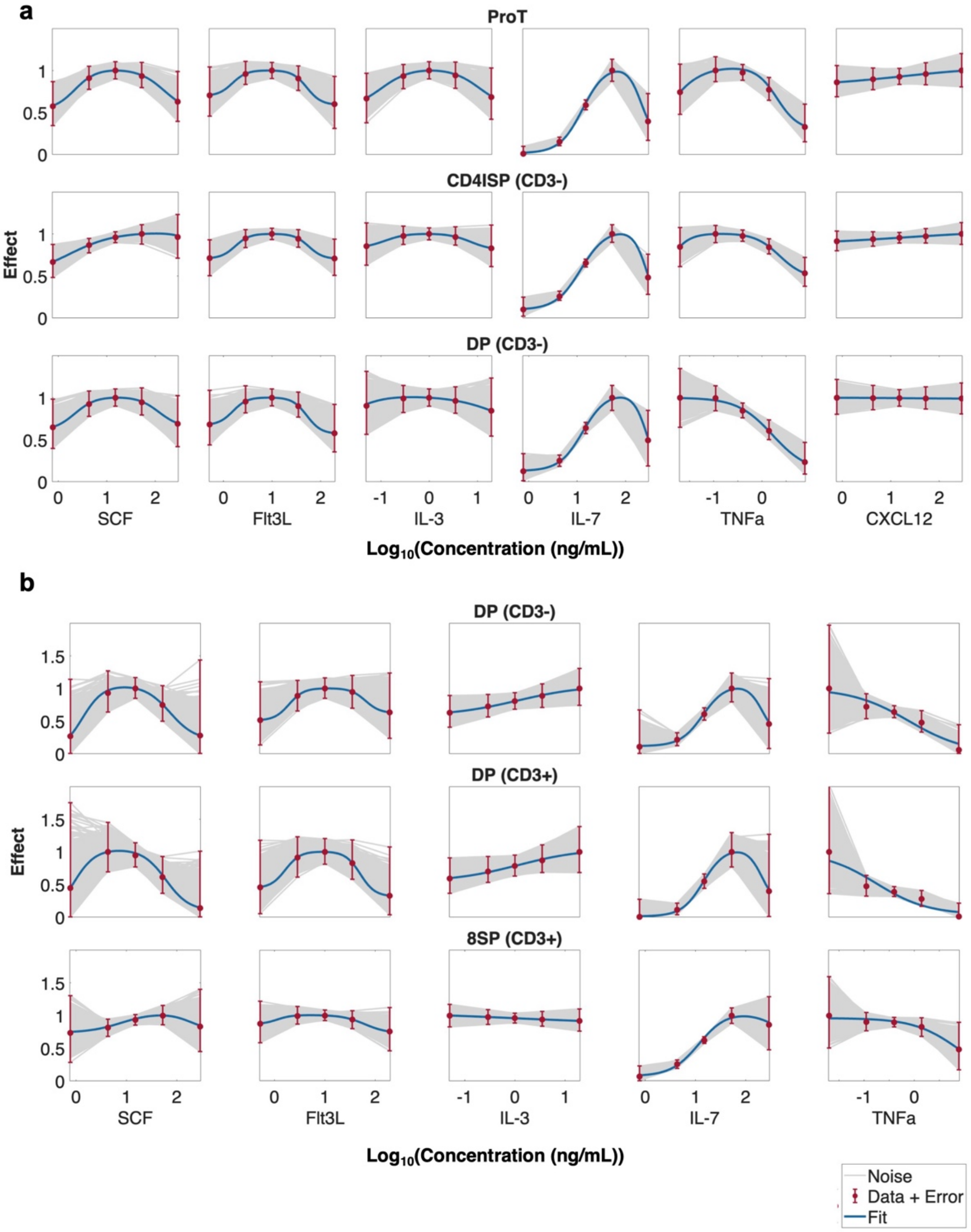


**Figure S6. Fits of single agent dose-response successfully describe non-monotonic behaviours. a)** Model predictions for cytokines SCF, Flt3L, IL-3, IL-7, TNFa and CXCL12 in cell differentiation (days 0-7 of the experiment) for ProT, CD4ISP and DP cells. **b)** Single fits for cytokines SCF, Flt3L, IL-3, IL-7 and TNFa in cell maturation (days 7-21 of the experiment) for DP and 8SP cells. **a-b)** Red dots and error bars: measurement ±2 standard deviations. Blue solid lines: model prediction. Grey lines: generated samples within ±2 standard deviations of experimental data (see Methods).

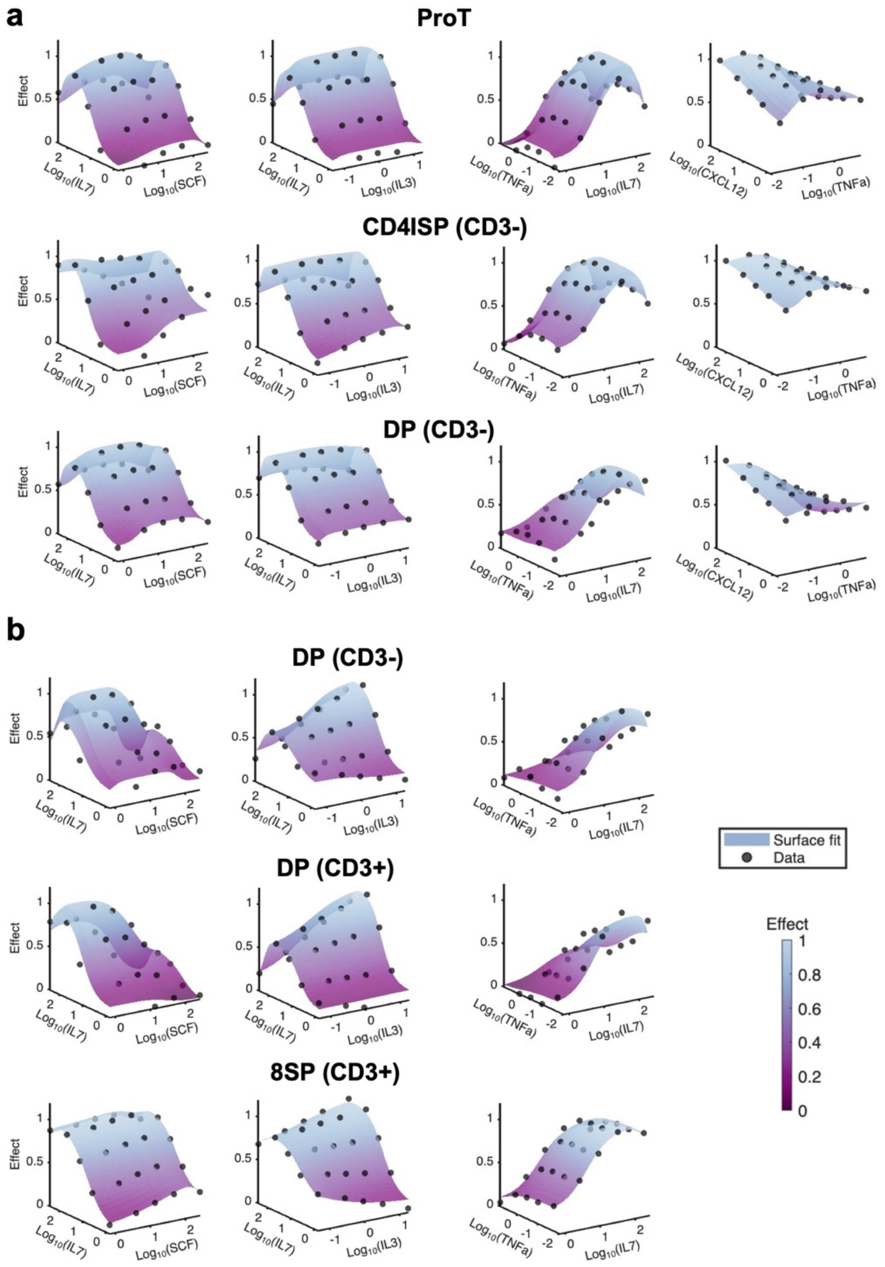


**Figure S7. Combination surface fits successfully recapitulate non-monotonic behaviours. a)** Model predictions of cytokine combinations SCF-IL7, IL3-IL7, IL7-TNFa and TNFa-CXCL12 in cell differentiation (days 0-7 of the experiment) for ProT, CD4ISP and DP cells. **b)** Model predictions of cytokine combinations SCF-IL7, IL3-IL7 and IL7-TNFa in cell maturation (days 7-21 of the experiment) for DP and 8SP cells. **a-b)** Black dots: data generated from polynomial regression coefficients. Scaled purple surfaces: combination fits with our new model. The axes for the cytokine concentrations are the physical concentrations from the previous experiment[10]. The vertical axis is the effect (between 0 and 1).

| | | | INTERACTION | | | |
|---|---|---|---|---|---|---|
| | | | **SCF/ IL7** | **IL3/ IL7** | **IL7/ TNFa** | **TNFa/ CXCL12** |
| *Days 7-14* | **CELL TYPE** | **Pro-T** | | | | CXCL12 (mono) |
| | | **CD4ISP** | | | | CXCL12 (mono) |
| | | **CD3$^-$** | | | | TNFa (mono) & CXCL12 (mono) |
| *Days 14-28* | | **CD3$^-$** | | IL3 (mono) | TNFa (mono) | |
| | | **CD3$^+$** | | IL3 (mono) | TNFa (mono) | |
| | | **8SP** | | IL3 (mono) | TNFa (mono) | |

**Table S2. Combinations for which at least one agent was set to be monotonic.** Green denotes interactions for which both agents were set to be biphasic. Red denotes interactions in which at least one agent was set to be monotonic. Each red shaded cell indicates which agent in the interaction exhibited a different behaviour than what was observed in the single fits. Yellow shaded cells indicate an interaction that was not present for the corresponding cell types. The only interaction for which the behaviour was set to be different than what was observed is the TNFa/CXCL12 interaction in CD3- cells for days 7-14, where TNFa was set to be monotonic to obtain a better fit.

| | | | INTERACTION | | | |
|---|---|---|---|---|---|---|
| | | | **SCF/ IL7** | **IL3/ IL7** | **IL7/ TNFa** | **TNFa/ CXCL12** |
| *Days 7-14* | **CELL TYPE** | **Pro-T** | | | | |
| | | **CD4ISP** | SCF (mono) | IL3 (mono) | | (TNFa mono) |
| | | **CD3$^-$** | | IL3 (mono) | TNFa (mono) | (TNFa mono) |
| *Days 14-28* | | **CD3$^-$** | | IL3 (mono decreasing) | TNFa (multi) | |
| | | **CD3$^+$** | SCF (mono) | IL3 (mono increasing) | | |
| | | **8SP** | | IL3 (mono decreasing) | | |

**Table S3. Comparison of behaviour in single dose-responses vs. in interactions.** Green denotes interactions for which both agents exhibited the same behaviour in single fits and in their interaction. Red denotes interactions in which an agent exhibited a different behaviour, sometimes only for certain concentrations, when in the presence of the other agent. Each red shaded cell indicates which agent in the interaction exhibited a different behaviour than what was observed in the single fits. Grey shaded cells indicate an interaction that was not present for the corresponding cell types.

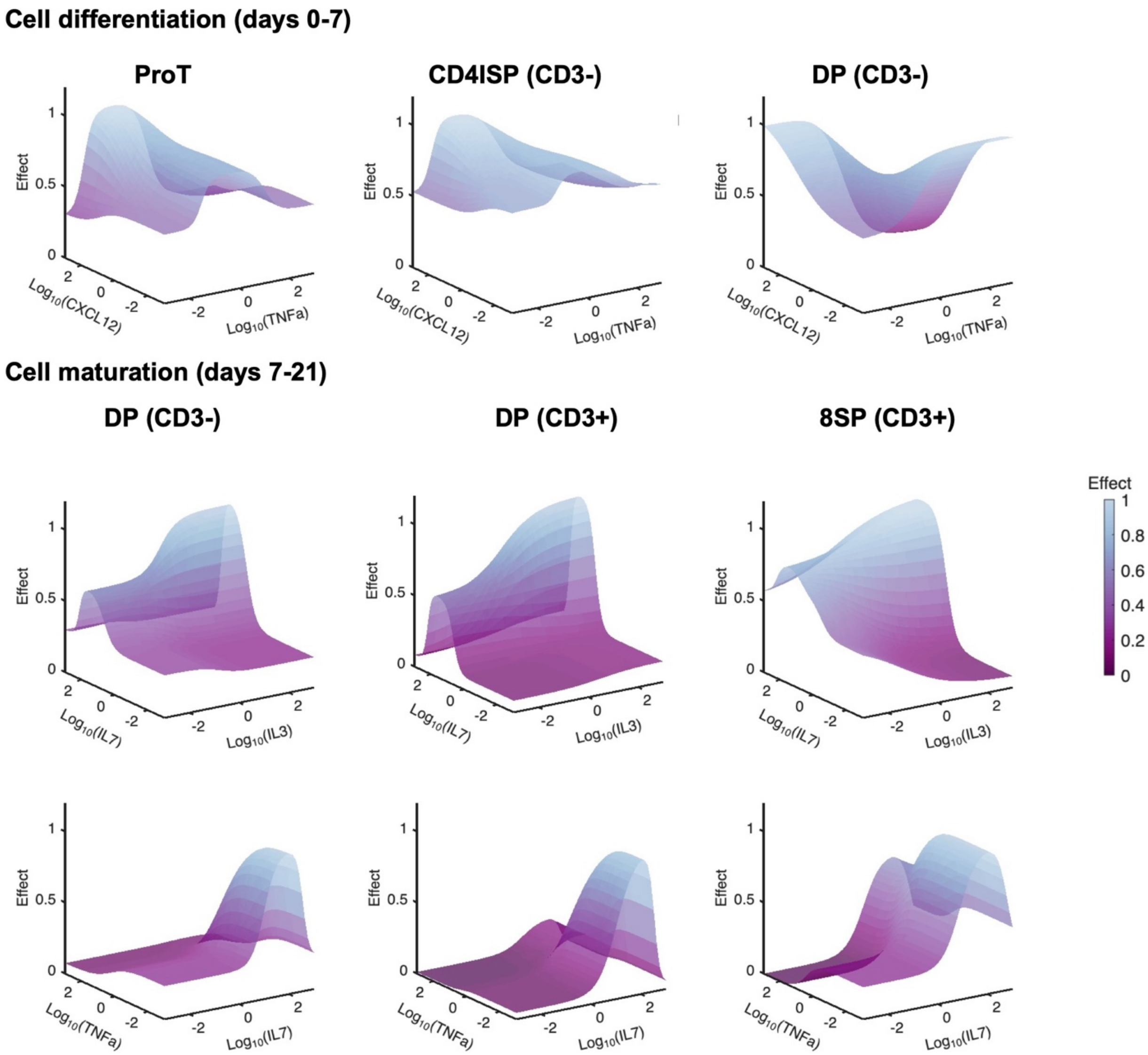


**Figure S8. Dose-response surfaces that included monotonic dose-responses.** All the surfaces for which at least one agent was monotonic. The surfaces were evaluated at extended concentrations values to show how the surfaces go back to the initial effect at higher concentrations.

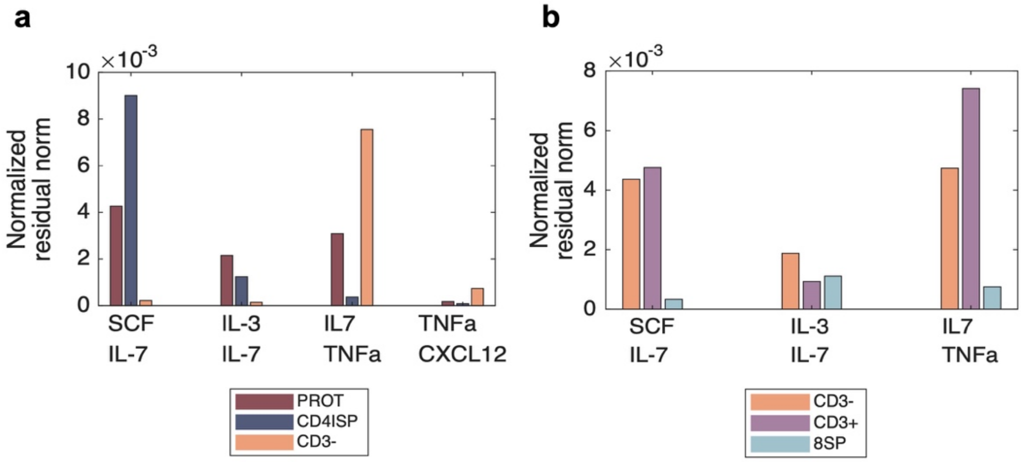


**Figure S9. Normalized residual norm of surface fits. a)** Normalized residual norm for the four interactions present in days 0-7. Red, dark purple and orange bars denote respectively ProT, CD4ISP and CD3- cells. **b)** Normalized residual norm (obtained by dividing the residual norm of the fit by the number of data points) for the three interactions present in days 7-21. **a-b)** Orange, purple and blue bars denote respectively CD3-, CD3+ and 8SP cells.

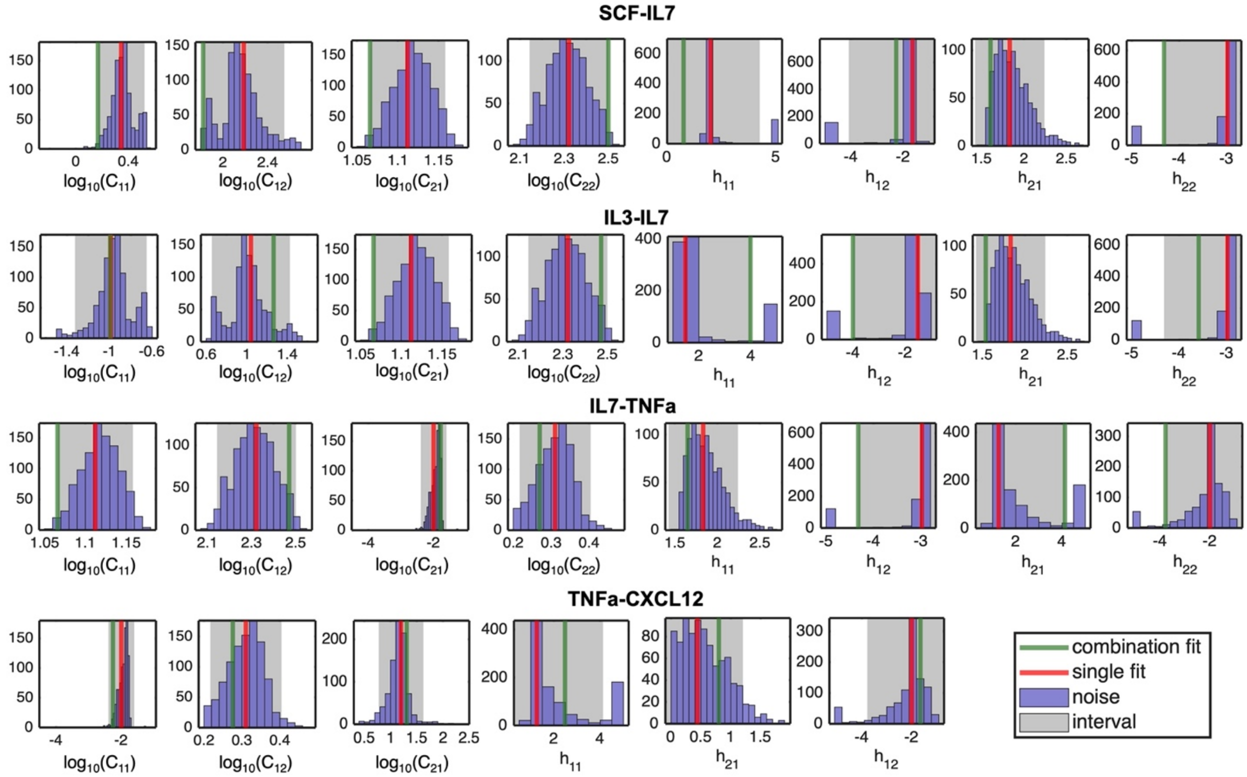


**Figure S10. Noise distributions for single agents in differentiation for ProT cells.** Noise distributions of the single agents for the ProT cells in the differentiation stage (days 0-7) in comparison with the single fits and the combination fits for EC50 and Hill slope parameters. Green: Combination fit estimate. Red: Single fit estimate. Purple: Distribution of noise for single agents. Grey: 95% confidence interval for the noise distributions.

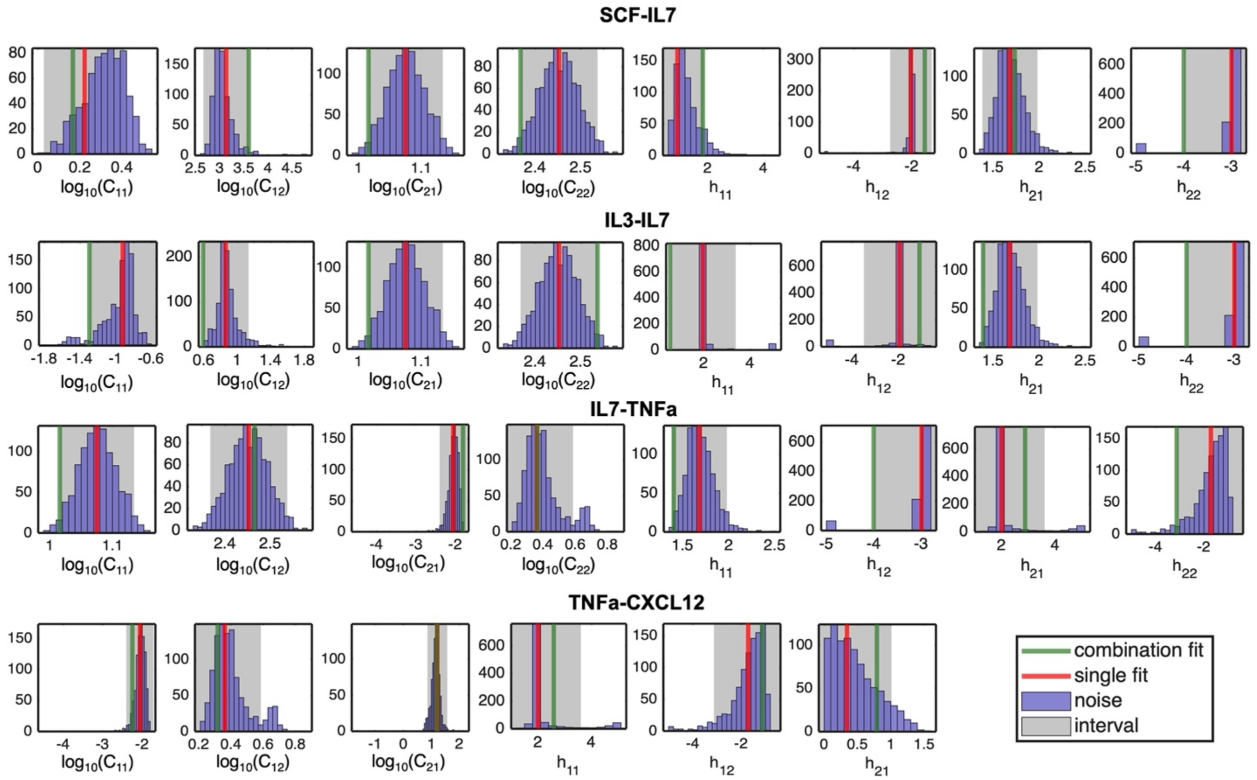


**Figure S11. Noise distributions for single agents in differentiation for C4ISP cells.** Noise distributions of the single agents for the CD4ISP cells in the differentiation stage (days 0-7) in comparison with the single fits and the combination fits for EC50 and Hill slope parameters. Green: Combination fit estimate. Red: Single fit estimate. Purple: Distribution of noise for single agents. Grey: 95% confidence interval for the noise distributions.

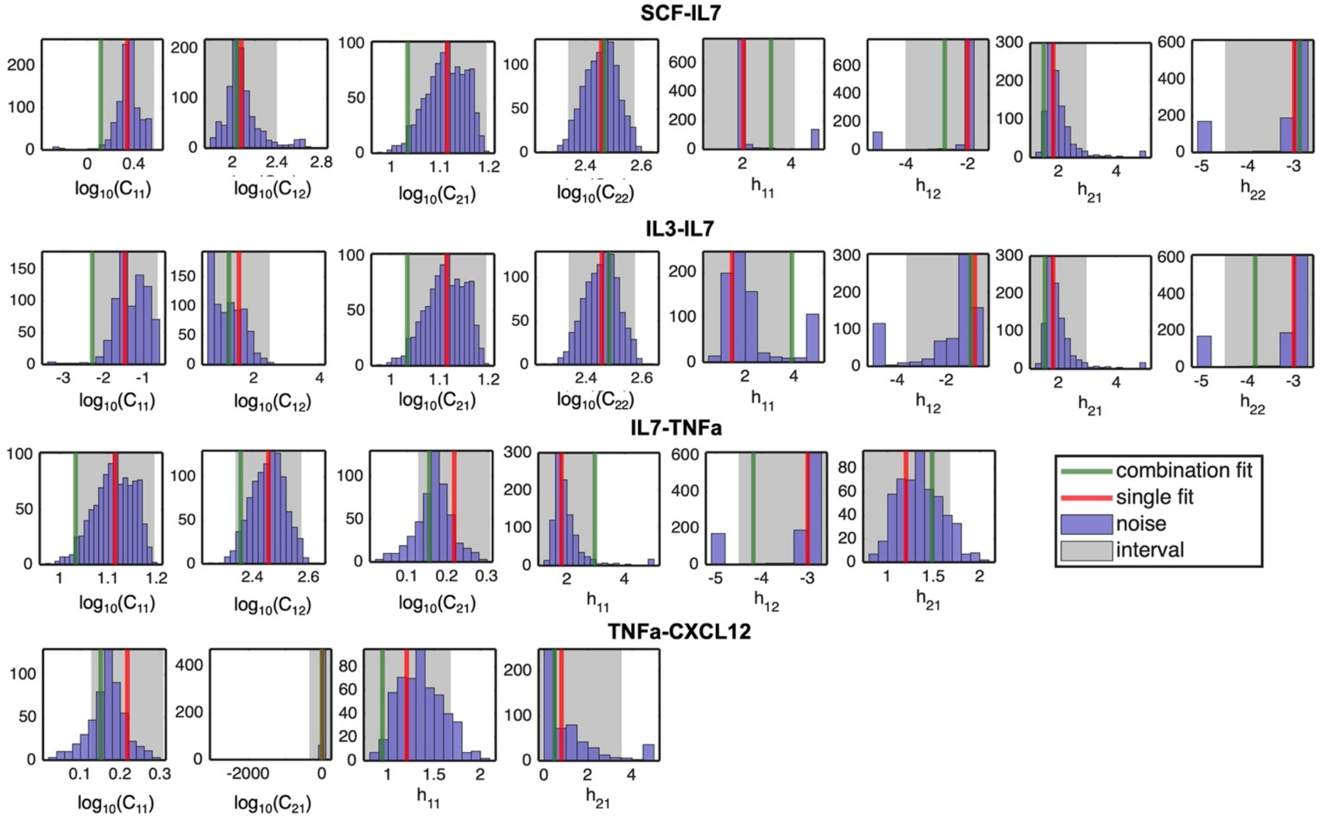


**Figure S12. Noise distributions for single agents in differentiation for CD3N cells.** Noise distributions of the single agents for the CD3N cells in the differentiation stage (days 0-7) in comparison with the single fits and the combination fits for EC50 and Hill slope parameters. Green: Combination fit estimate. Red: Single fit estimate. Purple: Distribution of noise for single agents. Grey: 95% confidence interval for the noise distributions.

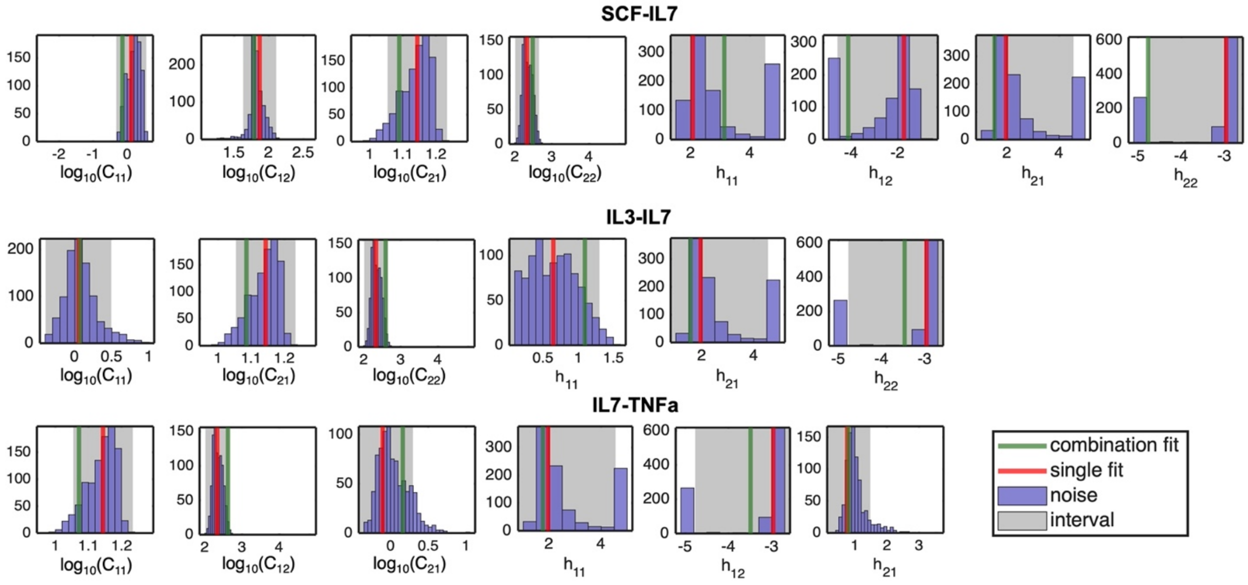


**Figure S13. Noise distributions for single agents in maturation for CD3N cells.** Noise distributions of the single agents for the CD3N cells in the maturation stage (days 7-21) in comparison with the single fits and the combination fits for EC50 and Hill slope parameters. Green: Combination fit estimate. Red: Single fit estimate. Purple: Distribution of noise for single agents. Grey: 95% confidence interval for the noise distributions.

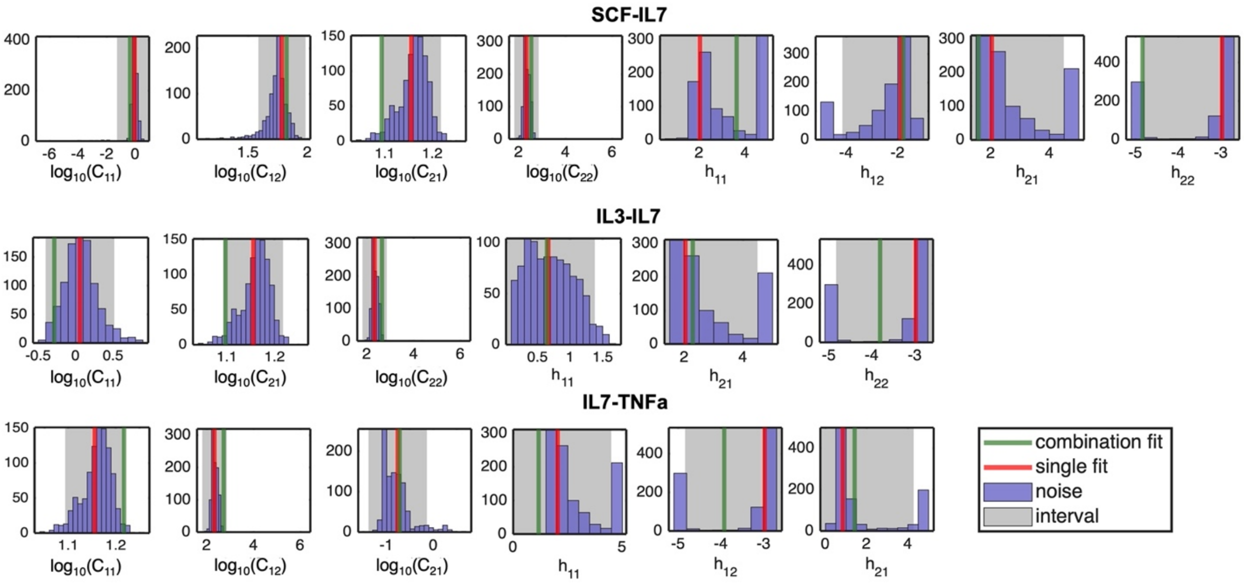


**Figure S14. Noise distributions for single agents in maturation for CD3P cells.** Noise distributions of the single agents for the CD3P cells in the maturation stage (days 7-21) in comparison with the single fits and the combination fits for EC50 and Hill slope parameters. Green: Combination fit estimate. Red: Single fit estimate. Purple: Distribution of noise for single agents. Grey: 95% confidence interval for the noise distributions.

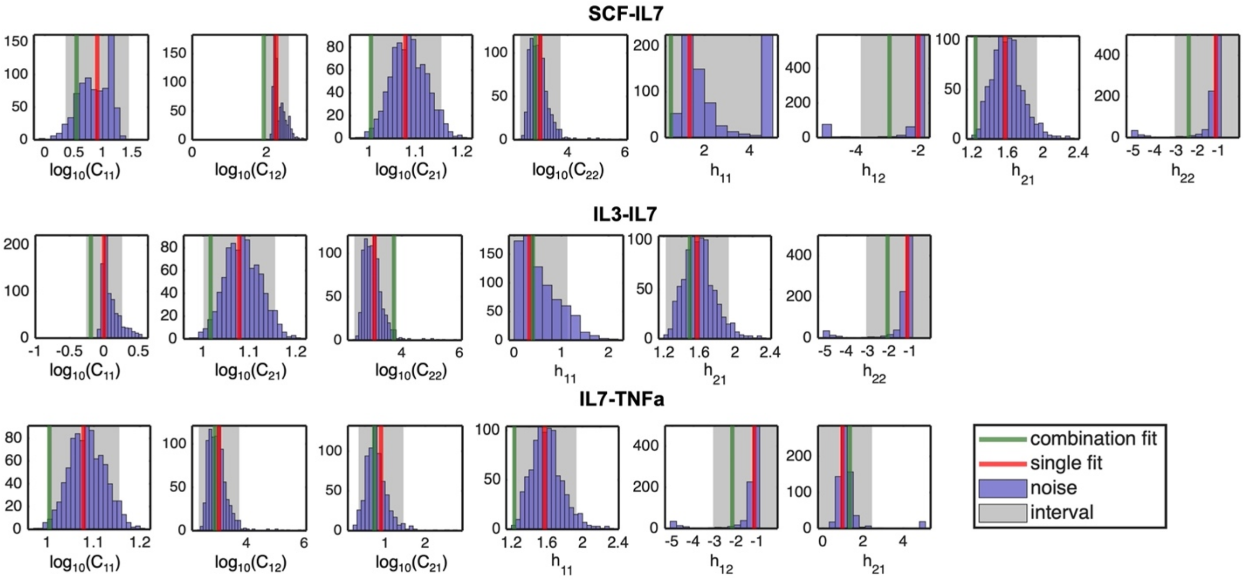


**Figure S15. Noise distributions for single agents in maturation for 8SP cells.** Noise distributions of the single agents for the 8SP cells in the maturation stage (days 7-21) in comparison with the single fits and the combination fits for EC50 and Hill slope parameters. Green: Combination fit estimate. Red: Single fit estimate. Purple: Distribution of noise for single agents. Grey: 95% confidence interval for the noise distributions.

### 6. Synergy parameter distributions

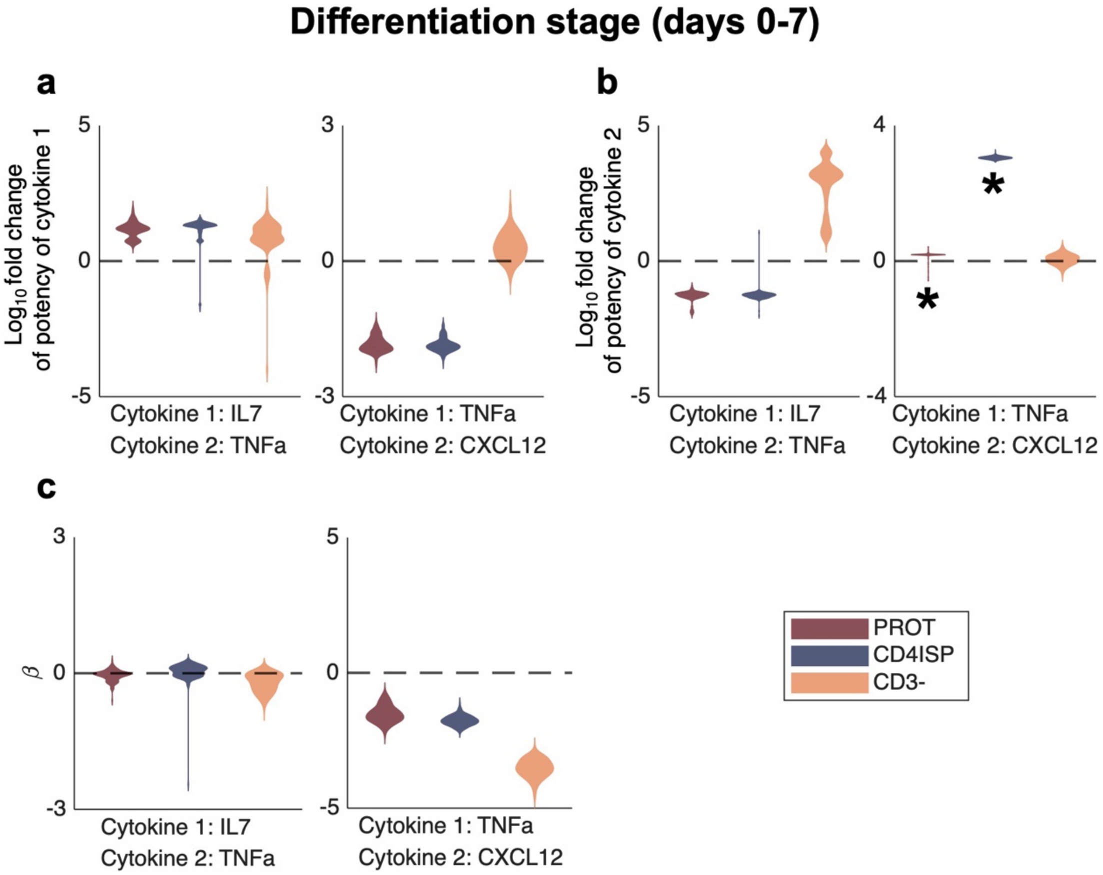


**Figure S16. Distributions of potency and efficacy parameters in cytokine combinations during differentiation (days 0-7) following Monte Carlo simulations. a)** Distributions of the fold change in potency of the first cytokine due to the presence of the second cytokine for combinations IL7-TNFa and TNFa-CXCL12 in days 0-7 of the experiment. **b)** Distributions of the fold change in potency of the second cytokine due to the presence of the first cytokine for combinations IL7-TNFa and TNFa-CXCL12 in days 0-7 of the experiment. **c)** Distributions of synergy of efficacy parameters ($\beta$) for cytokine combinations IL7-TNFa and TNFa-CXCL12 in days 0-7 of the experiment. **a-c)** First and second cytokines for each combination are shown on the x-axis (first cytokine on top and second cytokine below). Dotted line: no synergy of potency/efficacy. Above and below the dotted line denote synergistic and antagonistic potency/efficacy respectively. Black asterisk: distributions in which above the dotted line denotes antagonistic potency and below zero denotes synergistic potency due to the decreasing behaviour of the curve.

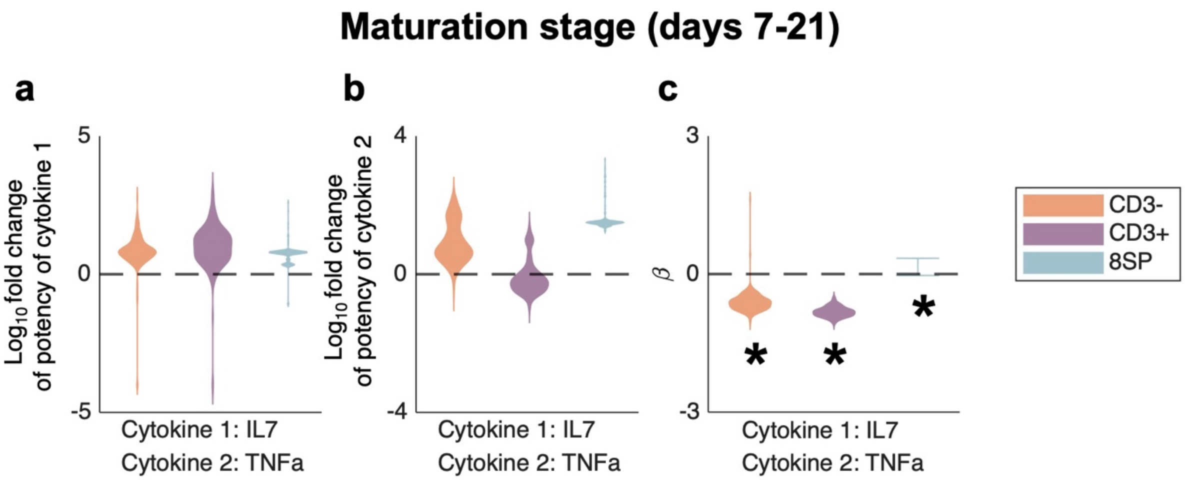


**Figure S17. Distributions of potency and efficacy parameters in cytokine combinations during maturation (days 7-21) following Monte Carlo simulations. a)** Distributions of the fold change in potency of the first cytokine due to the presence of the second cytokine for combination IL7-TNFa in days 7-21 of the experiment. **b)** Distributions of the fold change in potency of the second cytokine due to the presence of the first cytokine for combination IL7-TNFa in days 7-21 of the experiment. **c)** Distributions of synergy of efficacy parameters ($\beta$) for combination IL7-TNFa in days 7-21 of the experiment. **a-c)** First and second cytokines for each combination are shown on the x-axis (first cytokine on top and second cytokine below). Dotted line: no synergy of potency/efficacy. Above and below the dotted line denote synergistic and antagonistic potency/efficacy respectively. Black asterisk: distributions in which above the dotted line denotes antagonistic potency and below zero denotes synergistic potency due to the decreasing behaviour of the curve.

### 7. Parameter values

| | | CYTOKINES | | | | | |
|---|---|---|---|---|---|---|---|
| | | **SCF** | **FLT3L** | **IL3** | **IL7** | **TNFa** | **CXCL12** |
| **PARAMETERS AND CONFIDENCE INTERVALS** | $E_{0,1}$ | 0.52 [0.30, 0.74] | 0.69 [0.41, 0.98] | 0.54 [0.26, 0.83] | 0.01 [-0.02, 0.05] | 0.00 [-0.15, 0.15] | 0.81 [0.67, 0.95] |
| | $C_1$ | 0.35 [0.17, 0.52] | 0.19 [-0.14, 0.51] | -0.99 [-1.33, -0.65] | 1.11 [1.07, 1.16] | -2.00 [-2.39, -1.60] | 1.20 [0.78, 1.63] |
| | $h_1$ | 2.00 [-0.26, 4.26] | 3.00 [1.57, 4.42] | 1.48 [-1.03, 4.00] | 1.83 [1.43, 2.24] | 1.26 [-1.60, 4.12] | 0.44 [-0.32, 1.30] |
| | $E_{max1}$ | 1.01 [0.94, 1,08] | 1.00 [0.93, 1.08] | 1.02 [0.94, 1.09] | 1.05 [1.03, 1.07] | 1.05 [0.98, 1.12] | 1.05 [1.02, 1.08] |
| | $E_{0,2}$ | 0.49 [0.23, 0.75] | 0.59 [0.38, 0.80] | 0.54 [0.24, 0.85] | 0.16 [-0.05, 0.36] | 0.28 [0.10, 0.45] | |
| | $C_2$ | 2.19 [1.82, 2.55] | 1.72 [1.62, 1.81] | 1.04 [0.66, 1.43] | 2.32 [2.14, 2.50] | 0.31 [0.22, 0.40] | |
| | $h_2$ | -1.64 [-4.08, 0.80] | -2.94 [-4.56, -1.32] | -1.51 [-4.00, 0.98] | -2.98 [-4.32, -1.64] | -1.99 [-3.78, -0.20] | |
| | $E_{max2}$ | 1.01 [0.94, 1.08] | 1.00 [0.93, 1.08] | 1.01 [0.94, 1.09] | 1.01 [0.95, 1.08] | 1.00 [0.92, 1.08] | |

**Table S4. Single fit parameters and confidence interval bounds during cell differentiation (days 0-7) for ProT cells.** Eight parameters are fit for biphasic dose-responses. Grey cells denote parameters that were not fit because the curve was monotonic. Each fit parameter is shown with its 95% confidence interval in brackets [] using the generated noisy data. The $C_i$ parameters are in $\log_{10}$(concentration (ng/mL)).

| PARAMETERS AND CONFIDENCE INTERVALS | CYTOKINES | | | | | |
|---|---|---|---|---|---|---|
| | SCF | FLT3L | IL3 | IL7 | TNFa | CXCL12 |
| $E_{0,1}$ | 0.49 [0.31, 0.68] | 0.71 [0.55, 0.87] | 0.83 [0.59, 1.06] | 0.10 [0.02, 0.17] | 0.02 [-0.06, 0.09] | 0.86 [0.76, 0.97] |
| $C_1$ | 0.22 [0.03, 0.41] | 0.24 [0.13, 0.36] | -0.92 [-1.29, -0.55] | 1.08 [1.02, 1.13] | -2.06 [-2.40, -1.71] | 1.21 [0.86, 1.56] |
| $h_1$ | 0.93 [-0.01, 1.87] | 2.99 [2.01, 3.97] | 1.96 [0.55, 3.37] | 1.68 [1.38, 1.98] | 2.01 [0.41, 3.60] | 0.34 [-0.33, 1.01] |
| $E_{max1}$ | 1.02 [0.95, 1.09] | 1.00 [0.94, 1.06] | 1.00 [0.94, 1.06] | 1.05 [1.03, 1.07] | 1.01 [0.94, 1.07] | 1.05 [1.03, 1.07] |
| $E_{0,2}$ | 0.03 [-0.06, 0.12] | 0.70 [0.54, 0.86] | 0.80 [0.62, 0.99] | 0.00 [0.00, 0.00] | 0.47 [0.24, 0.70] | |
| $C_2$ | 3.14 [2.66, 3.61] | 1.76 [1.65, 1.86] | 0.86 [0.59, 1.13] | 2.45 [2.37, 2.54] | 0.36 [0.14, 0.59] | |
| $h_2$ | -2.04 [-2.75, -1.32] | -2.93 [-3.93, -1.93] | -1.95 [-3.49, -0.41] | -3.00 [-4.01, -1.99] | -1.71 [-3.13, -0.30] | |
| $E_{max2}$ | 1.00 [0.94, 1.07] | 1.00 [0.94, 1.06] | 1.00 [0.94, 1.06] | 1.01 [0.95, 1.07] | 1.00 [0.94, 1.07] | |

**Table S5. Single fit parameters and confidence interval bounds during cell differentiation (days 0-7) for CD4ISP cells.** Eight parameters are fit for biphasic dose-responses. Grey cells denote parameters that were not fit because the curve was monotonic. Each fit parameter is shown with its 95% confidence interval in brackets [] using the generated noisy data. The $C_i$ parameters are in $\log_{10}$(concentration (ng/mL)).

| PARAMETERS AND CONFIDENCE INTERVALS | CYTOKINES | | | | | |
|---|---|---|---|---|---|---|
| | SCF | FLT3L | IL3 | IL7 | TNFa | CXCL12 |
| $E_{0,1}$ | 0.60 [0.36, 0.85] | 0.67 [0.39, 0.95] | 0.73 [0.35, 1.11] | 0.13 [0.00, 0.25] | 1.00 [0.99, 1.01] | 1.00 [0.99, 1.01] |
| $C_1$ | 0.34 [0.11, 0.57] | 0.19 [-0.11, 0.50] | -1.47 [-2.29, -0.65] | 1.11 [1.03, 1.20] | 0.22 [0.13, 0.31] | 1.32 [-348.21, 350.84] |
| $h_1$ | 2.02 [-0.07, 4.12] | 3.00 [1.59, 4.40] | 1.46 [-0.99, 3.90] | 1.81 [0.65, 2.96] | 1.20 [0.72, 1.68] | 0.79 [-1.95, 3.52] |
| $E_{max1}$ | 1.01 [0.93, 1.08] | 1.00 [0.92, 1.08] | 1.00 [0.92, 1.08] | 1.05 [1.00, 1.10] | 0.11 [-0.04, 0.27] | 0.99 [0.83, 1.14] |
| $E_{0,2}$ | 0.64 [0.34, 0.93] | 0.57 [0.35, 0.78] | 0.54 [0.16, 0.93] | 0.00 [0.00, 0.00] | | |
| $C_2$ | 2.08 [1.76, 2.40] | 1.72 [1.60, 1.83] | 1.55 [0.62, 2.48] | 2.46 [2.34, 2.57] | | |
| $h_2$ | -2.01 [-4.04, 0.02] | -2.94 [-4.51, -1.36] | -0.84 [-3.65, 1.96] | -3.00 [-4.50, -1.49] | | |
| $E_{max2}$ | 1.01 [0.93, 1.08] | 1.00 [0.93, 1.08] | 1.02 [0.95, 1.10] | 1.01 [0.92, 1.09] | | |

**Table S6. Single fit parameters and confidence interval bounds during cell differentiation (days 0-7) for DP (CD3-) cells.** Eight parameters are fit for biphasic dose-responses. Grey cells denote parameters that were not fit because the curve was monotonic. Each fit parameter is shown with its 95% confidence interval in brackets [] using the generated noisy data. The $C_i$ parameters are in $\log_{10}$(concentration (ng/mL)).

| | | CYTOKINES | | | | |
|---|---|---|---|---|---|---|
| | | SCF | FLT3L | IL3 | IL7 | TNFa |
| PARAMETERS AND CONFIDENCE INTERVALS | $E_{0,1}$ | 0.02 [-0.30, 0.34] | 0.51 [0.16, 0.86] | 0.58 [0.41, 0.75] | 0.11 [-0.07, 0.29] | 1.00 [0.76, 1.24] |
| | $C_1$ | 0.12 [-0.32, 0.56] | 0.29 [0.02, 0.56] | 0.05 [-0.39, 0.49] | 1.14 [1.05, 1.23] | -0.10 [-0.50, 0.30] |
| | $h_1$ | 2.07 [-0.37, 4.52] | 2.99 [0.95, 5.03] | 0.65 [0.00, 1.30] | 1.96 [-0.64, 4.55] | 0.76 [0.04, 1.48] |
| | $E_{max1}$ | 1.01 [0.90, 1.12] | 1.00 [0.90, 1.11] | 1.05 [1.01, 1.09] | 1.05 [0.95, 1.15] | 0.00 [-0.04, 0.04] |
| | $E_{0,2}$ | 0.21 [-0.23, 0.64] | 0.62 [0.19, 1.05] | | 0.23 [-0.11, 0.56] | |
| | $C_2$ | 1.87 [1.63, 2.10] | 1.81 [1.24, 2.37] | | 2.33 [2.02, 2.65] | |
| | $h_2$ | -1.74 [-4.61, 1.13] | -2.95 [-4.72, -1.18] | | -2.96 [-4.77, -1.16] | |
| | $E_{max2}$ | 1.05 [0.97, 1.13] | 1.00 [0.89, 1.11] | | 1.01 [0.91, 1.12] | |

**Table S7. Single fit parameters and confidence interval bounds during cell maturation (days 7-21) for DP (CD3-) cells.** Eight parameters are fit for biphasic dose-responses. Grey cells denote parameters that were not fit because the curve was monotonic. Each fit parameter is shown with its 95% confidence interval in brackets [] using the generated noisy data. The $C_i$ parameters are in $log_{10}$(concentration (ng/mL)).

| PARAMETERS AND CONFIDENCE INTERVALS | CYTOKINES: SCF | FLT3L | IL3 | IL7 | TNFa |
|---|---|---|---|---|---|
| $E_{0,1}$ | 0.00 [-0.27, 0.28] | 0.44 [0.04, 0.84] | 0.54 [0.33, 0.75] | 0.01 [-0.06, 0.08] | 1.00 [0.73, 1.27] |
| $C_1$ | -0.05 [-1.29, 1.19] | 0.22 [-0.22, 0.65] | 0.06 [-0.40, 0.52] | 1.16 [1.09, 1.22] | -0.76 [-1.39, -0.14] |
| $h_1$ | 2.03 [-0.62, 4.67] | 2.99 [1.18, 4.81] | 0.67 [-0.04, 1.38] | 2.04 [-0.41, 4.49] | 0.84 [-2.60, 4.28] |
| $E_{max1}$ | 1.04 [0.90, 1.18] | 1.00 [0.87, 1.14] | 1.05 [0.99, 1.11] | 1.05 [0.94, 1.16] | 0.04 [-0.15, 0.23] |
| $E_{0,2}$ | 0.09 [-0.28, 0.46] | 0.31 [-0.01, 0.64] | | 0.16 [-0.17, 0.49] | |
| $C_2$ | 1.78 [1.58, 1.98] | 1.70 [1.46, 1.94] | | 2.32 [1.80, 2.84] | |
| $h_2$ | -1.88 [-4.16, 0.39] | -2.96 [-4.97, -0.95] | | -2.98 [-4.84, -1.12] | |
| $E_{max2}$ | 1.01 [0.88, 1.14] | 1.01 [0.87, 1.14] | | 1.01 [0.89, 1.14] | |

**Table S8. Single fit parameters and confidence interval bounds during cell maturation (days 7-21) for DP (CD3+) cells.** Eight parameters are fit for biphasic dose-responses. Grey cells denote parameters that were not fit because the curve was monotonic. Each fit parameter is shown with its 95% confidence interval in brackets [] using the generated noisy data. The $C_i$ parameters are in $\log_{10}$ (concentration (ng/mL)).

| | | CYTOKINES | | | | |
|---|---|---|---|---|---|---|
| | | SCF | FLT3L | IL3 | IL7 | TNFa |
| PARAMETERS AND CONFIDENCE INTERVALS | $E_{0,1}$ | 0.72 [0.41, 1.04] | 0.86 [0.53, 1.18] | 1.05 [1.01, 1.09] | 0.07 [-0.02, 0.16] | 0.96 [0.88, 1.04] |
| | $C_1$ | 0.91 [0.36, 1.46] | 0.05 [-0.40, 0.50] | 0.00 [-0.26, 0.26] | 1.08 [1.00, 1.16] | 0.90 [0.34, 1.46] |
| | $h_1$ | 1.35 [-1.88, 4.57] | 2.99 [1.77, 4.21] | 0.32 [-0.48, 1.13] | 1.58 [1.22, 1.93] | 0.97 [-0.50, 2.43] |
| | $E_{max1}$ | 1.02 [0.94, 1.11] | 1.00 [0.93, 1.07] | 0.87 [0.75, 0.98] | 1.05 [1.05, 1.05] | 0.00 [0.31, 0.31] |
| | $E_{0,2}$ | 0.75 [0.23, 1.17] | 0.73 [0.52, 0.94] | | 0.03 [-0.23, 0.29] | |
| | $C_2$ | 2.27 [1.93, 2.61] | 1.79 [1.59, 2.00] | | 3.05 [2.35, 3.75] | |
| | $h_2$ | -2.00 [-3.86, -0.14] | -1.94 [-4.03, 0.15] | | -1.15 [-3.08, 0.78] | |
| | $E_{max2}$ | 1.02 [0.95, 1.09] | 1.01 [0.94, 1.07] | | 1.03 [0.98, 1.08] | |

**Table S9. Single fit parameters and confidence interval bounds during cell maturation (days 7-21) for 8SP cells.** Eight parameters are fit for biphasic dose-responses. Grey cells denote parameters that were not fit because the curve was monotonic. Each fit parameter is shown with its 95% confidence interval in brackets [] using the generated noisy data. The $C_i$ parameters are in $\log_{10}$(concentration (ng/mL)).

| | | COMBINATIONS | | | |
|---|---|---|---|---|---|
| | | SCF-IL7 | IL3-IL7 | IL7-TNFa | TNFa-CXCL12 |
| PARAMETERS | $E_0$ | 1.05 | 1.05 | 1.05 | 1.02 |
| | $E_1$ | 0.90 | 0.91 | 0.16 | 0.29 |
| | $E_2$ | 0.23 | 0.15 | 0.72 | 0.82 |
| | $E_3$ | 0.00 | 0.08 | 0.00 | 0.48 |
| | $h_{11}$ | -0.76 | -3.98 | -1.65 | -2.48 |
| | $h_{12}$ | 2.27 | 4.00 | 4.32 | 1.64 |
| | $h_{21}$ | -1.61 | -1.54 | -4.12 | -0.80 |
| | $h_{22}$ | 4.32 | 3.59 | 3.78 | |
| | $C_{11}$ | 0.17 | -0.99 | 1.07 | -2.26 |
| | $C_{12}$ | 1.82 | 1.27 | 2.47 | 0.28 |
| | $C_{21}$ | 1.07 | 1.07 | -1.78 | 1.31 |
| | $C_{22}$ | 2.50 | 2.47 | 0.27 | |
| | $\alpha_{12}$ | 1.17 | 1.09 | 0.99 | 2.36 |
| | $\alpha_{21}$ | 0.31 | 1.65 | 1.08 | 0.18 |

**Table S10. Combinations fit parameters during cell differentiation (days 0-7) for ProT cells.** Grey cells denote parameters that were not fit because a cytokine exhibited a monotonic dose-response. The $C_i$ parameters are in $\log_{10}$(concentration (ng/mL)).

| | | COMBINATIONS | | | |
|---|---|---|---|---|---|
| | | SCF-IL7 | IL3-IL7 | IL7-TNFa | TNFa-CXCL12 |
| PARAMETERS | $E_0$ | 0.95 | 1.05 | 1.05 | 1.02 |
| | $E_1$ | 1.05 | 0.99 | 0.33 | 0.52 |
| | $E_2$ | 0.48 | 0.49 | 0.89 | 0.90 |
| | $E_3$ | 0.18 | 0.06 | 0.07 | 0.70 |
| | $h_{11}$ | -1.84 | -0.55 | -1.40 | -2.59 |
| | $h_{12}$ | 1.56 | 1.10 | 4.01 | 1.13 |
| | $h_{21}$ | -1.73 | -1.39 | -2.88 | -0.79 |
| | $h_{22}$ | 4.01 | 4.00 | 3.13 | |
| | $C_{11}$ | 0.17 | -1.29 | 1.02 | -2.25 |
| | $C_{12}$ | 3.60 | 0.59 | 2.46 | 0.32 |
| | $C_{21}$ | 1.02 | 1.02 | -1.81 | 1.20 |
| | $C_{22}$ | 2.37 | 2.54 | 0.37 | |
| | $\alpha_{12}$ | 0.62 | 1.22 | 1.30 | 1110.78 |
| | $\alpha_{21}$ | 0.14 | 0.09 | 0.99 | 0.09 |

**Table S11. Combinations fit parameters during cell differentiation (days 0-7) for CD4ISP cells.** Grey cells denote parameters that were not fit because a cytokine exhibited a monotonic dose-response. The $C_i$ parameters are in $\log_{10}$(concentration (ng/mL)).

| | | COMBINATIONS | | | |
|---|---|---|---|---|---|
| | | SCF-IL7 | IL3-IL7 | IL7-TNFa | TNFa-CXCL12 |
| PARAMETERS | $E_0$ | 1.05 | 1.05 | 0.00 | 0.00 |
| | $E_1$ | 0.91 | 0.87 | 0.00 | 1.05 |
| | $E_2$ | 0.34 | 0.34 | 0.96 | 1.05 |
| | $E_3$ | 0.16 | 0.13 | 0.48 | 0.51 |
| | $h_{11}$ | -3.15 | -3.90 | -2.96 | -0.94 |
| | $h_{12}$ | 2.76 | 1.01 | 4.18 | |
| | $h_{21}$ | -1.48 | -1.52 | -1.48 | -0.48 |
| | $h_{22}$ | 2.88 | 3.84 | | |
| | $C_{11}$ | 0.11 | -2.29 | 1.03 | 0.15 |
| | $C_{12}$ | 2.04 | 1.24 | 2.36 | |
| | $C_{21}$ | 1.03 | 1.03 | 0.16 | 0.24 |
| | $C_{22}$ | 2.47 | 2.48 | | |
| | $\alpha_{12}$ | 1.02 | 1.24 | 1433.66 | 0.15 |
| | $\alpha_{21}$ | 0.43 | 0.12 | 0.77 | 7.38 |

**Table S12. Combinations fit parameters during cell differentiation (days 0-14) for DP (CD3-) cells.** Grey cells denote parameters that were not fit because a cytokine exhibited a monotonic dose-response. The $C_i$ parameters are in $\log_{10}$(concentration (ng/mL)).

| | | COMBINATIONS | | |
|---|---|---|---|---|
| | | SCF-IL7 | IL3-IL7 | IL7-TNFa |
| PARAMETERS | $E_0$ | 1.05 | 1.05 | 0.08 |
| | $E_1$ | 0.47 | 0.66 | 0.07 |
| | $E_2$ | 0.28 | 0.21 | 1.01 |
| | $E_3$ | 0.12 | 0.30 | 0.23 |
| | $h_{11}$ | -3.13 | -1.10 | -1.76 |
| | $h_{12}$ | 4.15 | | 3.49 |
| | $h_{21}$ | -1.51 | -1.57 | -0.80 |
| | $h_{22}$ | 4.77 | 3.47 | |
| | $C_{11}$ | -0.14 | 0.07 | 1.07 |
| | $C_{12}$ | 1.79 | | 2.62 |
| | $C_{21}$ | 1.09 | 1.09 | 0.16 |
| | $C_{22}$ | 2.49 | 2.59 | |
| | $\alpha_{12}$ | 1.84 | 2.79 | 0.80 |
| | $\alpha_{21}$ | 0.92 | 6.89 | 1.17 |

**Table S13. Combinations fit parameters during cell maturation (days 7-21) for DP (CD3-) cells.** Grey cells denote parameters that were not fit because a cytokine exhibited a monotonic dose-response. The $C_i$ parameters are in $\log_{10}$(concentration (ng/mL)).

| | | COMBINATIONS | | |
|---|---|---|---|---|
| | | SCF-IL7 | IL3-IL7 | IL7-TNFa |
| PARAMETERS | $E_0$ | 1.05 | 1.05 | 0.21 |
| | $E_1$ | 0.31 | 0.57 | 0.01 |
| | $E_2$ | 0.11 | 0.15 | 1.01 |
| | $E_3$ | 0.00 | 0.08 | 0.05 |
| | $h_{11}$ | -3.63 | -0.65 | -1.17 |
| | $h_{12}$ | 1.78 | | 3.92 |
| | $h_{21}$ | -1.59 | -2.29 | -1.41 |
| | $h_{22}$ | 4.84 | 3.81 | |
| | $C_{11}$ | -0.37 | -0.29 | 1.22 |
| | $C_{12}$ | 1.82 | | 2.72 |
| | $C_{21}$ | 1.09 | 1.10 | -0.72 |
| | $C_{22}$ | 2.53 | 2.64 | |
| | $\alpha_{12}$ | 2.81 | 2.71 | 0.09 |
| | $\alpha_{21}$ | 2.17 | 0.15 | 1.53 |

**Table S14. Combinations fit parameters during cell maturation (days 7-21) for DP (CD3+) cells.** Grey cells denote parameters that were not fit because a cytokine exhibited a monotonic dose-response. The $C_i$ parameters are in $\log_{10}$(concentration (ng/mL)).

| | | COMBINATIONS | | |
|---|---|---|---|---|
| | | SCF-IL7 | IL3-IL7 | IL7-TNFa |
| PARAMETERS | $E_0$ | 1.05 | 1.05 | 0.67 |
| | $E_1$ | 0.98 | 0.82 | 0.00 |
| | $E_2$ | 0.54 | 0.07 | 1.04 |
| | $E_3$ | 0.00 | 0.53 | 0.23 |
| | $h_{11}$ | -0.52 | -0.40 | -1.23 |
| | $h_{12}$ | 2.93 | | 2.18 |
| | $h_{21}$ | -1.24 | -1.49 | -1.33 |
| | $h_{22}$ | 2.42 | 2.08 | |
| | $C_{11}$ | 0.55 | -0.19 | 1.00 |
| | $C_{12}$ | 1.93 | | 2.92 |
| | $C_{21}$ | 1.00 | 1.02 | 0.73 |
| | $C_{22}$ | 2.89 | 3.73 | |
| | $\alpha_{12}$ | 1.45 | 32.38 | 1.72 |
| | $\alpha_{21}$ | 0.09 | 20.98 | 1.34 |

**Table S15. Combinations fit parameters during cell maturation (days 7-21) for 8SP cells.** Grey cells denote parameters that were not fit because a cytokine exhibited a monotonic dose-response. The $C_i$ parameters are in $\log_{10}$(concentration (ng/mL)).

| | | | Mean $\log(\alpha_{12})$ | Mean $\log(\alpha_{21})$ | Mean $\beta$ |
|---|---|---|---|---|---|
| COMBINATION AND CELL TYPE | SCF-IL7 | ProT | $0.72 \pm 0.85$ | $-0.56 \pm 0.07$ | $-0.02 \pm 0.21$ |
| | | CD4ISP | $-0.25 \pm 0.38$ | $-0.69 \pm 0.13$ | $-0.13 \pm 0.12$ |
| | | DP (CD3-) | $0.36 \pm 0.06$ | $-0.53 \pm 0.03$ | $0.11 \pm 0.05$ |
| | IL3-IL7 | ProT | $1.37 \pm 1.46$ | $-0.55 \pm 0.07$ | $0.06 \pm 0.14$ |
| | | CD4ISP | $0.06 \pm 0.73$ | $-0.36 \pm 0.06$ | $-0.07 \pm 0.09$ |
| | | DP (CD3-) | $1.30 \pm 0.88$ | $-0.73 \pm 0.25$ | $-0.15 \pm 0.15$ |
| | IL7-TNFa | ProT | $-1.29 \pm 0.20$ | $1.15 \pm 0.29$ | $-0.05 \pm 0.12$ |
| | | CD4ISP | $-1.26 \pm 0.19$ | $1.18 \pm 0.50$ | $0.04 \pm 0.21$ |
| | | DP (CD3-) | $2.77 \pm 0.96$ | $0.83 \pm 0.75$ | $-0.21 \pm 0.20$ |
| | TNFa-CXCL12 | ProT | $0.17 \pm 0.12$ | $-1.84 \pm 0.18$ | $-1.52 \pm 0.31$ |
| | | CD4ISP | $3.05 \pm 0.01$ | $-1.86 \pm 0.15$ | $-1.76 \pm 0.18$ |
| | | DP (CD3-) | $0.05 \pm 0.15$ | $0.32 \pm 0.29$ | $-3.53 \pm 0.34$ |

**Table S16. Distribution means and standard deviations for potency and efficacy parameters during cell differentiation (days 0-7).** Means are shown with their standard deviations (mean ± standard deviation).

| | | | Mean $\log(\alpha_{12})$ | Mean $\log(\alpha_{21})$ | Mean $\beta$ |
|---|---|---|---|---|---|
| COMBINATION AND CELL TYPE | SCF-IL7 | DP (CD3-) | 0.56 ± 0.39 | -0.62 ± 0.16 | 0.11 ± 0.31 |
| | | DP (CD3+) | 0.91 ± 0.72 | -0.41 ± 0.77 | -0.29 ± 0.44 |
| | | 8SP | 0.08 ± 0.15 | -0.09 ± 0.03 | 0.01 ± 0.03 |
| | IL3-IL7 | DP (CD3-) | 0.71 ± 0.39 | 0.85 ± 0.06 | 1.19 ± 0.33 |
| | | DP (CD3+) | 1.28 ± 0.40 | -0.73 ± 0.29 | 1.51 ± 0.56 |
| | | 8SP | 2.81 ± 0.66 | 1.03 ± 0.53 | 0.36 ± 0.20 |
| | IL7-TNFa | DP (CD3-) | 0.91 ± 0.57 | 0.82 ± 0.66 | -0.61 ± 0.23 |
| | | DP (CD3+) | -0.09 ± 0.49 | 0.99 ± 0.81 | -0.82 ± 0.11 |
| | | 8SP | 1.59 ± 0.36 | 0.71 ± 0.37 | 0.02 ± 0.05 |

**Table S17. Distribution means and standard deviations for potency and efficacy parameters during cell maturation (days 7-21).** Means are shown with their standard deviations (mean ± standard deviation).
